# A more comprehensive and reliable analysis of individual differences with generalized random forest for high-dimensional data: validation and guidelines

**DOI:** 10.1101/2025.10.28.685232

**Authors:** Jinwoo Lee, Junghoon Justin Park, Maria Pak, Seung Yun Choi, Jiook Cha

## Abstract

Analyzing individual differences in treatment or exposure effects is a central challenge in psychology and behavioral sciences. Conventional statistical models have focused on average treatment effects, overlooking individual variability, and struggling to identify key moderators. Generalized Random Forest (GRF) can predict individualized treatment effects, but current implementations suffer from two critical limitations: (1) prediction performances vary substantially across random initializations, and (2) identification of key moderator is limited in high-dimensional settings. Here, we introduce two methodological advances to address these issues. First, a seed ensemble strategy stabilizes predictions by aggregating models trained under different random initializations. Second, a backward elimination procedure systematically identifies key moderators from high-dimensional inputs. Simulation analyses across diverse scenarios demonstrate that our approach achieves reliable and valid predictions across random seeds, improved performance in moderator identification, and robust generalization to independent data. To facilitate adoption and interpretation, we provide step-by-step guidance using large-scale neuroimaging dataset (*N* = 8,778) with reusable code. These enhancements make GRF more reliable for modeling individual differences in treatment effects, supporting data-driven hypothesis generation, and identification of responsive subgroups.

## Introduction

*Why are some children more vulnerable to stress after early-life traumatic event?* (Belsky & Pluess, 2009; Binder et al., 2008; Caspi et al., 2003; Shonkoff et al., 2012) *Why do some older adults maintain cognitive function despite neurodegenerative risk?* (Ngandu et al., 2015; Solomon et al., 2018; Stern, 2012) *Why is antidepressant response heterogeneous across individuals?* (Bousman et al., 2017; Greden et al., 2019; Rush et al., 2006)

These questions highlight a central issue in psychology and the behavioral sciences: identifying how and for whom a treatment or exposure exerts its effects, and the key moderators of those effects. Understanding individual differences in treatment response is crucial not only for refining theoretical models of mental processes but also for translating research into clinical and practical applications. In particular, it enables identification of subgroups who most likely to benefit from an intervention, thereby advancing personalized and precision medicine (Bryan et al., 2021; Feuerriegel et al., 2024).

Conventional statistical approaches have long been used to model such variability. However, despite notable contributions, they face fundamental challenges. For instance, univariate moderation analyses, which test the significance of an interaction between an outcome and a single moderator, may overlook more potent sources of individual differences that are not explicitly modeled (Memon et al., 2019). Even when higher-order interaction terms are included, regression models require researchers to pre-specify interactions, reducing power and increasing risk of biased interpretation (Akimova et al., 2021; Kent et al., 2018). Furthermore, their reliance on linear models limits their ability to capture complex interplay among moderators, especially in the high-dimensional settings of modern neuroscience where countless variables could plausibly influence a treatment effect (Figlio et al., 2017; Fu et al., 2024; Warrier et al., 2021). For example, a conventional linear regression analysis aiming to account for all interactions among 45 covariates would require examining over 35 trillion interaction terms (Cha et al., 2024). Exhaustively evaluating such interactions is both computationally infeasible and statistically unsound.

Generalized Random Forest (GRF) offers a powerful alternative for studying individual differences through non-parametric and data-driven approach (Athey et al., 2019). GRF provides estimates of the average treatment effect (ATE) as well as individualized treatment effects (ITEs), and it identifies moderators’ nonlinear combinations underlying heterogeneity in treatment effects. This enables researchers to uncover and interpret treatment effect moderators without pre-specifying interaction terms or conducting targeted hypothesis tests. This flexibility makes GRF particularly valuable for informing personalized decision-making and generating novel hypotheses centered around empirically identified moderators.

Despite its promise, current applications of the GRF framework in the behavioral sciences face two critical limitations. First, GRF models exhibit substantial variability in ITE prediction performance depending on the random seed used during training, raising concerns about the reliability of both the model and its outputs. Second, although GRF can handle more moderators than regression models, it is substantially underpowered when modeling ITEs involving the extensive set of potential moderators typically reported in contemporary psychology and the behavioral sciences — for example, whole-brain distributed circuits involved in moderating stress responses from threatening input (Bo et al., 2024; Cha et al., 2014; Sinha et al., 2016; Wang et al., 2022). These issues highlight the need for a more reliable and comprehensive analytical framework that can identify robust moderators in high-dimensional spaces and remain robust to stochastic variation like random initialization.

In this paper, we first provide a brief overview of how GRF works and how it has been applied in psychology and behavioral sciences. We then present two simulation-based methodological proposals designed to enhance the reliability and comprehensiveness of GRF-based analyses. Finally, we demonstrate a step-by-step tutorial of our proposed framework on a real-world dataset, covering model fitting, model selection, ATE and ITE evaluation, and the interpretation of key moderators.

## Brief Overview of GRF

GRF is a causal machine learning technique that computes ATE and ITEs based on the potential outcome framework. By stratifying samples based on their predicted ITEs, each sample can be characterized as belonging to high- or low-response group regardless of their treatment or exposure status and key moderators can be identified (see **Fig 1**). Since all individual difference analyses are based on the ITEs, it is essential to understand the internal mechanism by which GRF computes these quantities. In this section, we first review how ITEs are typically defined and calculated in conventional causal inference frameworks and then outline how GRF improves upon these methods. We then summarize how GRF has been applied to generate novel insights in psychology and the behavioral sciences.

**Fig 1.**
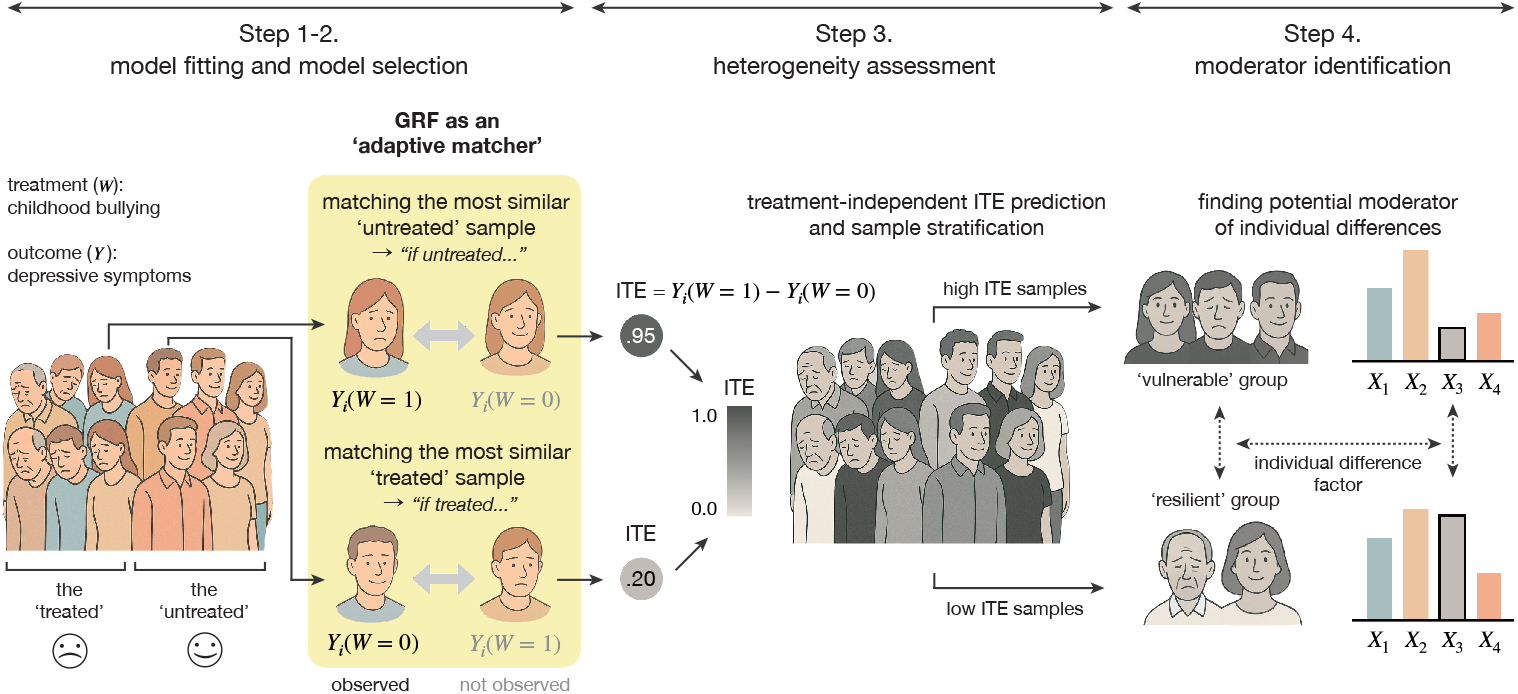
Conceptual scheme of GRF-based individual differences analyses. GRF acts as an ‘adaptive matcher’ to predict an ITE for a given sample *i* by identifying similar samples with different treatment values based on covariate profiles. It assigns weights to these neighboring samples according to their similarity in covariates and uses the weighted outcomes to calculate the sample *i*’s counterfactual outcome value. Applying this procedure across all samples allows for out-of-bag prediction of ITEs regardless of treatment assignment (see **Box 1**). Based on the predicted ITEs, samples are stratified by treatment group to identify covariates closely associated with individual variability in ITEs. The background illustration was generated by ChatGPT-4o.

### ITE calculation as a matching problem

It is helpful to start with the potential outcome framework (Rubin, 2005), a theoretical background of GRF in causal inference. In this framework, the individual treatment effect for sample *i*, τ_*i*_, is defined as follows:

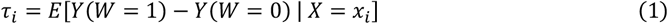

where *Y* and *W* represent the observed outcome and treatment values, respectively. Also, *x*_*i*_ is the covariate^**1**^ set for sample *i*. However, we cannot observe both potential outcomes (*W* = 1 and *W* = 0) for the same individual. Only one is realized, a limitation known as the “fundamental problem of causal inference” (Holland, 1986). Here, two causal assumptions, unconfoundedness and overlap, can address this issue. Unconfoundedness assumes that treatment assignment is based only on observed variables (Rubin, 1978), and overlap ensures that every unit has a positive chance of receiving each treatment (Rosenbaum & Rubin, 1983). When these assumptions are met, equation (1) can be mathematically rewritten in the following form that is identifiable in observational studies (Rosenbaum & Rubin, 1983):

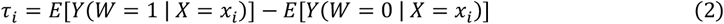

Hence, to calculate an ITE of sample *i*, samples that display the most similar covariate profile to need to be chosen from the treated and untreated samples (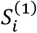 and 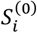, respectively). For example, one of the causal machine learning techniques, *k*-nearest neighborhood (*k*NN) matching, selects 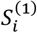 and 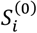 based on Euclidean distance in the covariate space (Abadie & Imbens, 2006). However, since this approach results in inaccurate matching in high-dimensional feature spaces (Beyer et al., 1999; Radovanović et al., 2010; Zimek et al., 2012), a more efficient algorithm is required to accurately estimate τ_*i*_ in settings with many moderator candidates.

### GRF as an adaptive matcher for calculating ITE

To improve the identification of 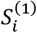 and 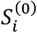 in high-dimensional settings, GRF acts like an adaptive matching machine that clusters individuals based on their covariate profiles related to variability of treatment effects regardless of how complex the rules might be.

Technically, GRF employs a greedy split algorithm of classical random forest. It builds trees by selecting random subsamples and splitting leaves to maximize the squared differences in leaf-level ATEs, effectively separating subsamples based on covariates profiles related to treatment effects. By assembling all trees, GRF assigns a similarity-based weight to each sample on how frequently it falls into the same leaf node as sample *i* to calculate τ_*i*_. These weights are then used to compute τ_*i*_ as a weighted average of the outcome values of the other samples. Unlike *k*NN matching, which relies on simple distance metrics in high-dimensional space, GRF adaptively learns complex matching rules through recursive splits, allowing it to identify locally relevant covariates and avoid the curse of dimensionality. Compared to the *k*NN matching, GRF recovered simulated ITEs with five times less mean-squared error (Athey et al., 2019; Wager & Athey, 2018).

Applying decision tree methods that partition samples into leaves based on specific variables, GRF can automatically model nonlinear interactions among moderators that drive variability in ITEs. Additionally, by tracking how frequently each candidate moderator is used to split the trees, the algorithm provides a measure of variable importance, offering a data-driven opportunity to identify key moderators that engage in treatment effect heterogeneity.

Also, GRF provides ITEs for every sample without explicit train/test data splitting, while preventing data leakage problem during model fitting and prediction of ITEs (Athey et al., 2019; Wager & Athey, 2018). For a detailed explanation of how GRF internally predicts each sample’s ITE in a robust manner, see **Box 1**.

### Application of GRF in psychology and the behavioral sciences

Given the complex and heterogeneous nature of human behavior, brain function, and their relationship, GRF is emerging as a valuable tool in psychology and the behavioral sciences. It allows for flexible, non-parametric computation of treatment effects that can account for a wide range of covariates, providing more personalized insights than traditional methods can offer.

GRF has been increasingly utilized in psychological studies, demonstrating its value in understanding diverse mental health outcomes. Notable research includes studies on suicide risk (Goldman-Mellor et al., 2022; Ross et al., 2024; Solomonov et al., 2023), trauma (Shiba et al., 2021, 2023, 2022), dementia (Nakagomi et al., 2025), and well-being (Komura et al., 2023). In particular, the recent study provides a compelling example of GRF’s utility in psychological research, specifically addressing a critical research gap concerning the heterogeneous effects of retirement on older adults’ mental health (Fu et al., 2024), a salient inquiry given global demographic shift.

Previous studies on the mental health consequences of retirement have produced inconsistent findings, often attributable to methodological heterogeneity or the failure to appropriately model individual differences in treatment effects. A major limitation of prior work lies in the widespread assumption of homogeneous treatment effects. By leveraging GRF’s high functional flexibility and its ability to identify relevant covariates in high-dimensional data, Fu and their colleagues effectively bypassed this limitation and provided crucial scientific insights: retirement generally improved mental health by reducing depression and increasing well-being, but notably, its depression-reducing effects were more pronounced for specific subgroups, such as overweight males, males aged 57-67, and males with internet habits. These findings highlight the value of GRF’s data-driven approach in uncovering subgroup-specific effects and identifying the covariates that contribute most to treatment heterogeneity. This allows for more targeted and personalized policy interventions, aimed at optimizing retirement-related mental health benefits and improving the well-being of older adults.

As summarized in **Table 1**, these studies showcase GRF’s capacity to handle large sample sizes (ranging from 1,150 to 196,610) and numerous covariates (from 7 to 128). The covariates in these studies predominantly include sociodemographic variables, psychosocial factors, and health-related factors. GRF’s ability to model nonlinear and multivariate relationships makes it ideal for capturing the intricate interplay of genetic, neural, and environmental factors that contribute to psychological outcomes.

**Table 1.** Recent applications of GRF in psychology and the behavioral sciences.

| Reference | Sample Size | The number of covariates | Treatment | Outcome | Covariates |
| --- | --- | --- | --- | --- | --- |
| Nakagomi et al., 2025 | 5,451 | 31 | Internet use | Dementia | • demographic characteristics<br>• socioeconomic status<br>• social relations<br>• health conditions<br>• functional capacity<br>• behavioral factor |
| Garrison-Desany et al., 2024 | 2,943 | 128 | PTSD | Substance use (tobacco, alcohol, cannabis) | • sociodemographic variables<br>• childhood trauma<br>• social interaction<br>• chronic stress<br>• mindfulness<br>• sleep quality<br>• area deprivation index<br>• etc |
| Ross et al., 2024 | 196,610 | not specified | Psychiatric hospitalization | Suicide attempts within 12 months | • psychopathologic risk factors<br>• physical disorders and treatments<br>• facility-level quality indicators<br>• indicators of social determinants of health |
| Fu et al., 2024 | 8,432 ~ 12,364 | 33 | Retirement | Depression<br>Subjective well-being | • marital status<br>• life satisfaction<br>• total work income<br>• job satisfaction<br>• personal relationships |
| Komura et al., 2023 | 16,164 | 37 | Life satisfaction | Cognitive functioning | • sociodemographic variables<br>• health-related factors<br>• psychological factors<br>• social factors<br>• personality traits |
| Solomonov et al., 2023 | 16,164 | 7 | Emotional support | Thoughts of suicide | • sociodemographic variables |
| Shiba et al., 2023 | 2,664 ~ 3,350 | 55 | Disaster damage (home loss) | Functional limitations (physical disability, activities of daily living, instrumental activities of daily living) | • sociodemographic variables<br>• health conditions<br>• psychosocial factors<br>• behavioral factors |
| Goldman-Mellor et al., 2022 | 57,312 | 39 | Hospitalization | Suicide within 30 days and 1 year | • patient-level characteristics<br>• hospital-level characteristics |
| Shiba et al., 2022 | 1,150 ~ 1,644 | 51 | Disaster damage (home loss, loss of loved ones) | Depressive symptoms, posttraumatic stress symptoms | • sociodemographic variables<br>• health conditions<br>• psychosocial factors<br>• behavioral factors |
| Shiba et al., 2021 | 2,664 ~ 3,350 | 51 | Disaster damage (home loss, loss of loved ones) | Cognitive disability | • sociodemographic variables<br>• health conditions<br>• psychosocial factors<br>• behavioral factors |

## Current Limitations and Proposed Suggestions

Despite the methodological strengths of GRF in accurately and robustly predicting ITEs even in high-dimensional feature spaces, this section highlights two major challenges of the current GRF framework that hinder reliable and comprehensive analysis. We then propose two simple yet effective solutions to address these issues and demonstrate through simulation studies that our proposals can meaningfully mitigate these limitations. All analysis codes used in this section are publicly available in our GitHub repository (https://github.com/jinw00-lee/GRFprotocol).

### Toward more reliable framework

#### Limited reliability in ITE prediction

In practice, GRF models often exhibit considerable instability in the prediction of ITEs, depending heavily on the random seed used during training. Even when models are trained on the same data with identical hyperparameters, the predicted ITEs can vary substantially across seeds. In some cases, the model may yield predictions that align well with observed treatment–outcome structure under one seed, while performing poorly under another.

More technically, this variability is observed in the ‘differential forest prediction’ coefficient (hereafter, β_*ITE*_) produced by the calibration test — a diagnostic tool for evaluating model fit in GRF (see **Box 2** for details of calibration test). In the calibration test for the trained GRF model, two metrics, β_*ATE*_ and β_*ITE*_, evaluate how well the predicted ATE and ITE align with the observed treatment-outcome relationships for each sample. When each metric is statistically significant at *P* < .05 and its estimate is close to 1, it indicates that the model closely explains the variance of the observed outcome variable (see **Box 2** for details of calibration test). However, across different seeds, β_*ITE*_ estimates and their *P*-values can fluctuate substantially, often failing to reach statistical significance. This instability may undermine confidence in the model’s results and their interpretability.

In a simulation experiment using a dataset of 1,000 samples with 149 covariates, treatment, and outcome (see **Appendix Methods 1** for data simulation process), the estimates and statistical significance of β_*ITE*_ varied widely depending on the choice of single random seed used for model training (see **Fig 2a**). While β_*ATE*_ estimates remained stable across all seeds, ITE predictions significantly explained the variance in observed outcomes well for only half of the seeds, while failing to do so for the other half. This makes it difficult to determine whether the 149 covariates used in the analysis truly capture key moderators of individual-level treatment effects and may lead to misleading conclusions if results from only a favorable seed are reported (see **Fig 2a**).

**Fig 2.**
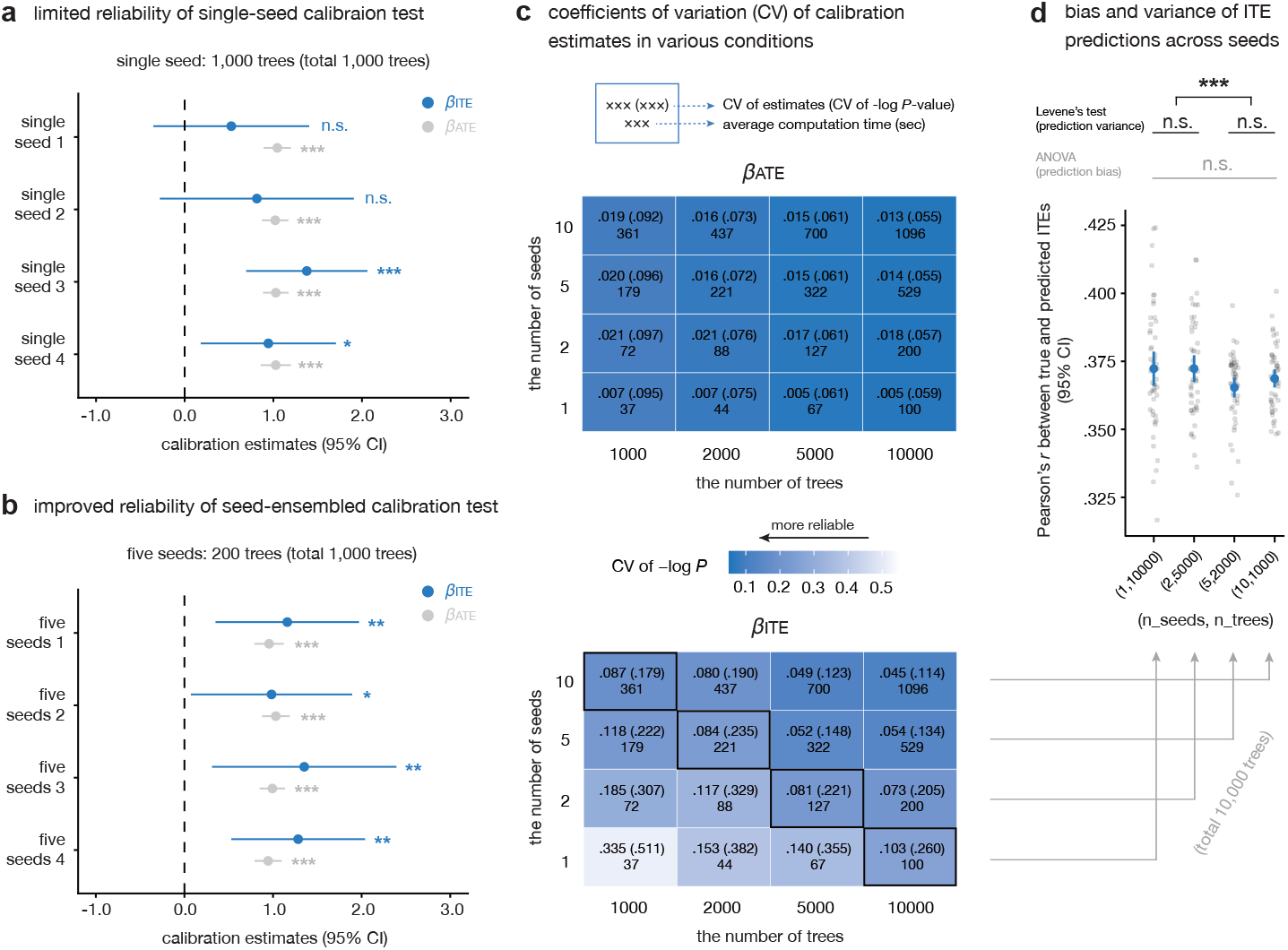
Reliability gains from seed-ensemble. **a**, Limited reliability of single-seed calibration test. A GRF model with 1,000 trees was fitted for each random seed, and calibration performance was evaluated. **b**, Improved reliability of seed-ensembled calibration test. For each seed condition, a “big model” was created by ensembling GRFs trained on five different random seeds, each with 200 trees (totaling 1,000 trees, as in panel a). **c**, Coefficients of variation (CV) of calibration estimates in various seed and tree conditions. CV was computed for estimated coefficients and their *P*-values across 50 repeated trials per each seed-tree condition. Darker blue indicates lower variability across trials (i.e., lower CV). This panel was visualized based on the simulation experiment with a nonlinear and weak heterogeneity condition. See **Appendix A** for results from other conditions (i.e., nonlinear-strong, linear-weak, linear-strong conditions). **d**, Bias and variance of ITE predictions across seeds. The distribution of ITE prediction performance across 50 trials is shown for four different conditions in the panel a, each using a total of 10,000 trees per model. The performance metric was calculated as the Pearson correlation between the simulated and the predicted ITEs. One-way ANOVA and Levene’s Test were applied to test statistical differences in mean and variance of prediction performance across conditions, respectively. All *P*-values were FDR-corrected. \**P*_FDR_ < .05; \*\**P*_FDR_ < .01; \*\*\**P*_FDR_ < .001

Such instability may arise from the inherent randomness in building random forest models (e.g., subsampling and random feature selection during tree growth) as well as from the substantial statistical uncertainty involved in ITE prediction itself (Athey et al., 2019).

#### Seed ensemble to improve reliability

As a practical solution to the instability of β_*ITE*_ estimates across random seeds, we propose the use of a *seed ensemble*. This approach involves training multiple GRF models using the same variables and hyperparameter configuration but with different random seeds and then aggregating them into a single ‘big model’. If each model contains *n* trees and *k* such models are combined, the resulting ensemble contains a total of *k × n* trees.

Importantly, we emphasize that constructing a model with *k × n* trees using a single seed is not equivalent to combining *k* models with *n* trees each using *k* different seeds. Although the total number of trees is the same, the ensemble trained across diverse seeds introduces greater structural variation and reduces individual sources of randomness. In our simulations, we evaluated the variability of calibration metrics under a seed-ensembled condition, where instead of training a GRF model with 1,000 trees using a single seed, we aggregated models trained with 200 trees each across five different seeds — totaling 1,000 trees. Unlike the single-seed condition in which calibration metrics varied greatly depending on the chosen seed (see **Fig 2a**), all big models under the seed-ensembled condition yielded β_*ITE*_ estimates that were statistically significant and similar in point estimates (see **Fig 2b**). These results suggest that, even with the same total number of trees, aggregating models trained under diverse seeds allows for a more reliable evaluation of calibration estimates given a fixed set of variables and hyperparameters.

We extended the simulation to examine how the variability of calibration estimates changes across different combinations of tree and seed counts. Specifically, we tested 16 conditions combining tree counts of 1,000, 2,000, 5,000, and 10,000 with seed counts of 1, 2, 5, and 10 (see **Fig 2c**). For each condition, we fitted 50 GRF models using different random seeds and analyzed how the coefficient of variation for β_*ATE*_ and β_*ITE*_ changed as a function of tree and seed combination (see **Appendix Methods 1**). We observed that coefficients of variation of the log-transformed *P*-values for β_*ITE*_ was lower in the seed ensembled condition compared to the single-seed condition (see **Fig 2c**), which implies reliability gain of calibration metrics. Interestingly, coefficients of variations of log-transformed *P*-values for β_*ATE*_ remained consistently low across all conditions, regardless of whether the seed ensemble was used (see **Fig 2c**). The results presented in **Fig 2c** are derived from conditions where ITEs are simulated under nonlinear function and weak heterogeneity, but we observed similar stabilizing effect of the seed ensemble on *β*_*ITE*_ across both linear and nonlinear treatment functions, as well as under varying degrees of treatment effect heterogeneity (see **Appendix A**).

However, it is important to note that using a seed ensemble required more computation time, even when the total number of trees was the same. For example, training a model with 10,000 trees under a single seed took approximately 100 seconds, whereas building a big model by aggregating GRF models with 1,000 trees each across 10 different seeds took about 360 seconds with eight CPU threads of MacBook Air M2-chip (run at up to 3.94 GHz per thread; see **Fig 2c**). Thus, while seed ensembling offers greater stability, it comes at a computational cost, highlighting the need to explore the most efficient combination by balancing stability and runtime.

In addition to examining a calibration metric of ITE prediction, we directly assessed the accuracy and reliability of ITE predictions across different seed and tree configurations. To control for the number of trees, we focused on four conditions from the original 16 conditions: (n_seeds, n_trees): [(1, 10,000), (2, 5,000), (5, 2,000), and (10, 1,000)], each yielding 10,000 trees per model. For each of the 50 models per condition, Pearson correlation between the true and predicted ITEs was computed as an accuracy metric. One-way ANOVA and Levene’s test were used to assess differences in the mean and variance of this metric across the four conditions (see **Appendix Methods 1** and **Fig 2d**).

While no significant differences in mean accuracy were observed across conditions (*F*(3, 196) = 1.951, *P* = .123), the variance differed significantly (*F*(3, 196) = 6.050, *P* = 6×10^-4^). Post-hoc tests revealed that models with five or ten seeds showed significantly lower variance in ITE prediction performance compared to those with only one or two seeds (*F*(1, 198) = 16.52, *P* = 7×10^-5^). This suggests that while seed ensembling does not reduce prediction bias, it substantially improves reliability. Considering the bias-variance trade-off in conventional machine learning, this highlights our seed ensemble as a promising strategy to enhance reliability without sacrificing bias.

Taken together, these findings suggest that the seed ensemble offers a more reliable strategy for ITE prediction, beyond the benefits obtained by simply increasing the number of trees under a fixed seed.

### Toward more comprehensive framework

#### Lack of comprehensive model selection framework

One of the key strengths of GRF over traditional linear models lies in its ability to model multifaceted interactions among massive covariates without requiring explicit specification of interaction terms. This makes GRF particularly effective for identifying treatment effect heterogeneity in a data-driven manner. Nevertheless, a critical limitation remains: the choice of which variables to include as optimal covariates must still be manually specified by the researcher. For instance, consider a study aiming to understand how bullying victimization in childhood influences children’s mental health problems, potentially moderated by neurobiological processing of such stressful events. Given that including too many covariates can underpower the GRF’s ability to estimate treatment effects (Athey & Wager, 2019; Sverdrup et al., 2025), there is a practical limit to the number of covariates that can be included in the model relative to sample size. As a result, researchers often rely on prior literature to hand-select a subset of variables known to modulate stress response such as amygdala volume (Ben-Zion et al., 2023), blood-oxygenation-level–dependent (BOLD) activities of the amygdala during specific tasks (Dick et al., 2021), or functional connectivity within the salience network (Zhang et al., 2022).

However, recent findings in neuroscience consistently suggest that the encoding of stressful events and the subsequent regulation is implemented in a distributed manner across the whole brain (Bo et al., 2024; Cha et al., 2014; Sinha et al., 2016; Wang et al., 2022). This raises the possibility that a small set of hand-picked variables may be suboptimal for capturing the full complexity of individual differences. In such cases, a more comprehensive modeling strategy would involve incorporating a much broader set of whole-brain features as covariates, allowing the GRF algorithm to identify the most relevant heterogeneity-factors in a data-driven fashion.

#### Backward elimination for comprehensive model selection

As a solution to the challenge of identifying the optimal subset of covariates from a large feature space in a data-driven manner, we propose a *backward elimination model selection* framework, which is designed to maximize the calibration of both ATE and ITE prediction with the observed dataset. This procedure consists of five steps: First, a full GRF model is trained using all *J* available covariates. A seed ensemble approach is used; whereby multiple models trained under different random seeds are merged to form a single ‘big model’. This initial model may display poor calibration performances due to its innate high-dimensionality. Second, the covariate with the lowest variable importance is removed. Third, this process is iterated until only one covariate remains, yielding a total of *J* big models. Fourth, we retain only those models that satisfy the following two criteria:

✓ **Calibration test:** Both β_*ATE*_ and β_*ITE*_ must be statistically significant at *P*_FDR_ < .05.
✓ **Overlap assumption:** All estimated propensity scores, *ê*, must lie within the range 0.05 ≤ *ê* < 0.95.

The calibration test ensures that the model satisfies a minimum standard for accurately predicting both ATE and ITEs (Athey & Wager, 2019), while the overlap assumption is a fundamental assumption for causal identification of treatment effects (see **Box 3** for conceptual details). Finally, among the subset of models that meet both criteria, we select the ‘best model’ based on the following fit index: fit index

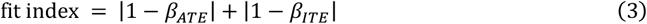

This metric summarizes how closely the model’s calibration coefficients align with their golden standard (i.e., 1), with lower values indicating that the predicted ATE and ITEs can generate the observed treatment-outcome relationship.

In simulation experiments, we confirmed that our backward elimination framework improved identification of the true effect-modifying factors from the full covariates. In this analysis, we evaluated whether the best model selected through backward elimination could accurately identify the three true key moderators among 149 input covariates (see **Appendix Methods 2**). Across both linear and nonlinear ITE generation conditions, our approach achieved higher F1 scores than the conventional heuristic that interprets the top 10% of variables by importance as key moderators (e.g., Athey & Wager, 2019; Gudmundsdottir et al., 2022; Shiba et al., 2021; see **Fig 3a**). This suggests that, in real-world datasets where true moderators are unknown, variables identified through backward elimination are more reliable than those derived from standard heuristics.

**Fig 3.**
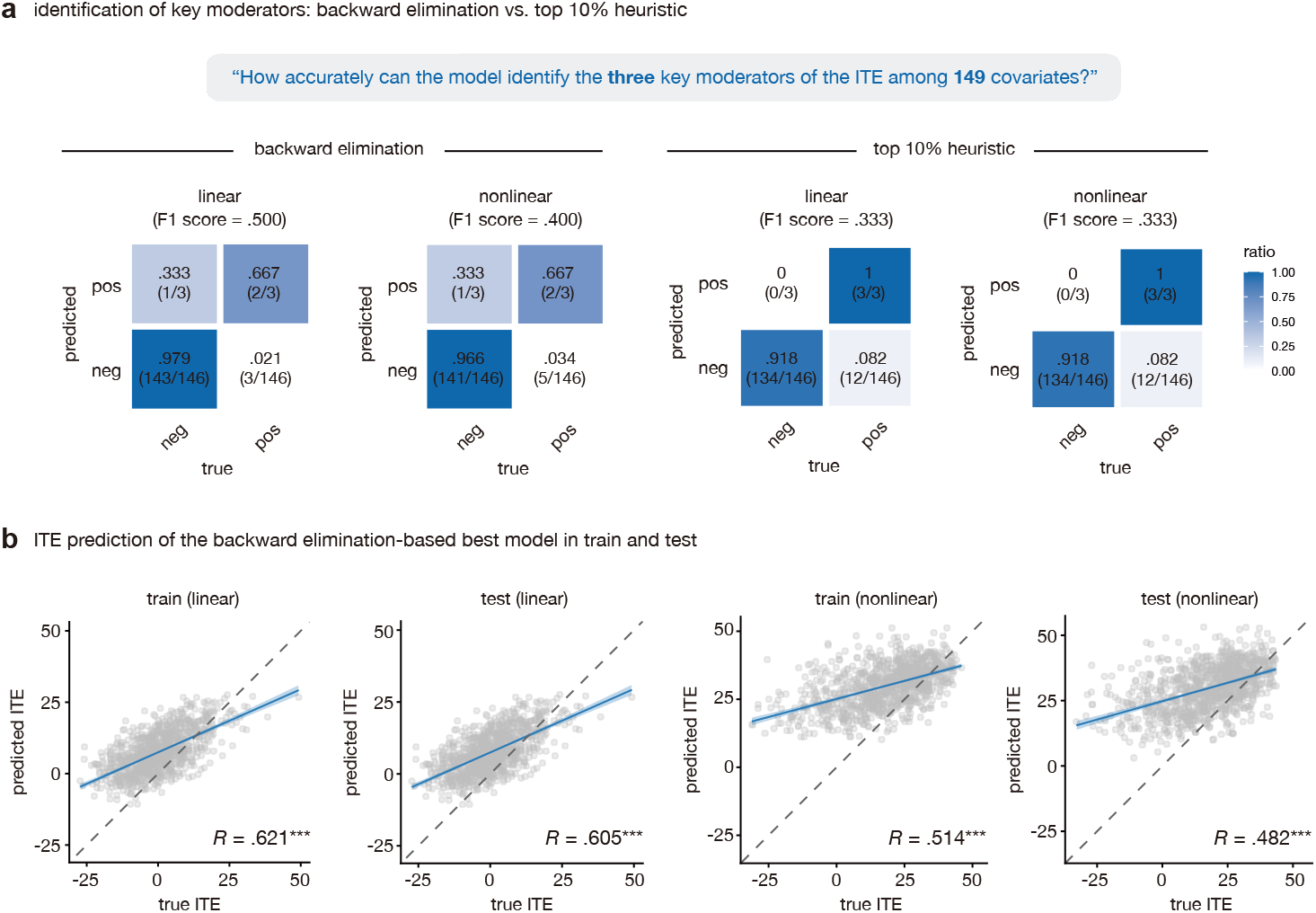
Accurate moderator identification and ITE predictions of backward-elimination model selection. **a**, Identification of key moderators: backward elimination vs. top 10% heuristic. We compared two methods for identifying true moderators in a simulation study where the ITE was generated from three out of 149 covariates. This comparison was conducted under conditions where the ITE was generated using linear and nonlinear functions, respectively (see **Appendix Methods 2**). F1 score was used as the performance metric. In each confusion matrix, the count and ratio of variables identified as key moderators and those that were not by each model are presented in the ‘pos’ and ‘neg’ columns, respectively. **b**, ITE prediction of the backward elimination-based best model in train and test. For each linear and nonlinear condition, we evaluated how accurately the best model identified through backward elimination predicted the ITE in both the train set (used for model selection) and the test set (not used). As a performance metric, the Pearson correlation between the true and predicted ITEs was computed. The test set shared the same ITE generation function as the train set but included noise in a different pattern (see **Appendix Methods 2**). The dotted diagonal line represents the identical function. \**P* < .05; \*\**P* < .01; \*\*\**P* < .001.

Furthermore, the best model selected via backward elimination showed comparable ITE prediction performance between the training and independent test sets. Although performance was generally higher under the linear condition than nonlinear one (linear: *R*_train_ = .621, *P*_train_ < 2×10^-16^, *R*_test_ = .605, *P*_test_ < 2×10^-16^; nonlinear: *R*_train_ = .514, *P*_train_ < 2×10^-16^, *R*_test_ = .482, *P*_test_ < 2×10^-16^), similar levels of predictive accuracy were observed across both conditions (see **Fig 3b**). Given that the test set shared the same ITE generation function as the training set but contained independently generated noise (see **Appendix Methods 2**), these results indicate that our backward elimination model selection captures the underlying ITE function robustly without overfitting to noise.

## Tutorial with Real-world Dataset

In this section, we demonstrate an automated analysis framework that incorporates seed ensemble and backward elimination–based model selection, using a real-world dataset. As an illustrative example, we assess the effects of childhood bullying exposure on depressive symptoms and examine individual differences in these effects. Additionally, we investigate which demographic, environmental, and neuroanatomical features are most strongly associated with these individual differences. All analysis codes used in the tutorial are publicly available in our GitHub repository (https://github.com/jinw00-lee/GRFprotocol) and **Appendix Codes**.

We applied this framework to baseline data from 8,778 children in the Adolescent Brain and Cognitive Development (ABCD) study (Casey et al., 2018; see **Appendix Methods 3** for details of data descriptions and preprocessing). The treatment variable was a binary variable `‘kbi_p_c_bully’`, which was derived from the Kiddie Schedule for Affective Disorder and Schizophrenia (K-SADS) interview (Kaufman et al., 2017), reflecting whether the child had experienced peer bullying (1 = “Yes”; 0 = “No”). The outcome variable was a continuous variable `‘cbcl_dsm_depression_t’` representing the child’s *t*-score of depressive symptoms from the Child Behavior Checklist (CBCL)(Achenbach, 2011).

A total of 138 covariates were assessed. These spanned demographics (i.e., age, biological sex, race/ethnicity, and caregivers’ age, marital status, income, and education), the rate of mental illness among family members (‘family disease ratio’)(Cha et al., 2024; Karcher et al., 2021; Rice et al., 1995), body mass index (‘BMI’)(Nishimi et al., 2022; Zhang et al., 2023; Zheng et al., 2024), and 84 gray matter volume measures across bilateral whole-brain regions. Among the sample, 7,544 children had no reported history of bullying, while 1,234 had experienced bullying (see **Appendix B** for demographics).

**Fig 4a** illustrates the causal model tested in this tutorial. In the GRF framework, the covariate set consists of variables that may confound both the cause and the outcome while also potentially moderating the treatment effect (see the footnote 1). While this simplified causal model is used for tutorial purposes, substantial effort is required in real research to validate the causal structure such as grounding it in prior studies or collecting covariates prior to the cause variable to block potential influences from the cause to covariates (see ‘Discussion’).

**Fig 4.**
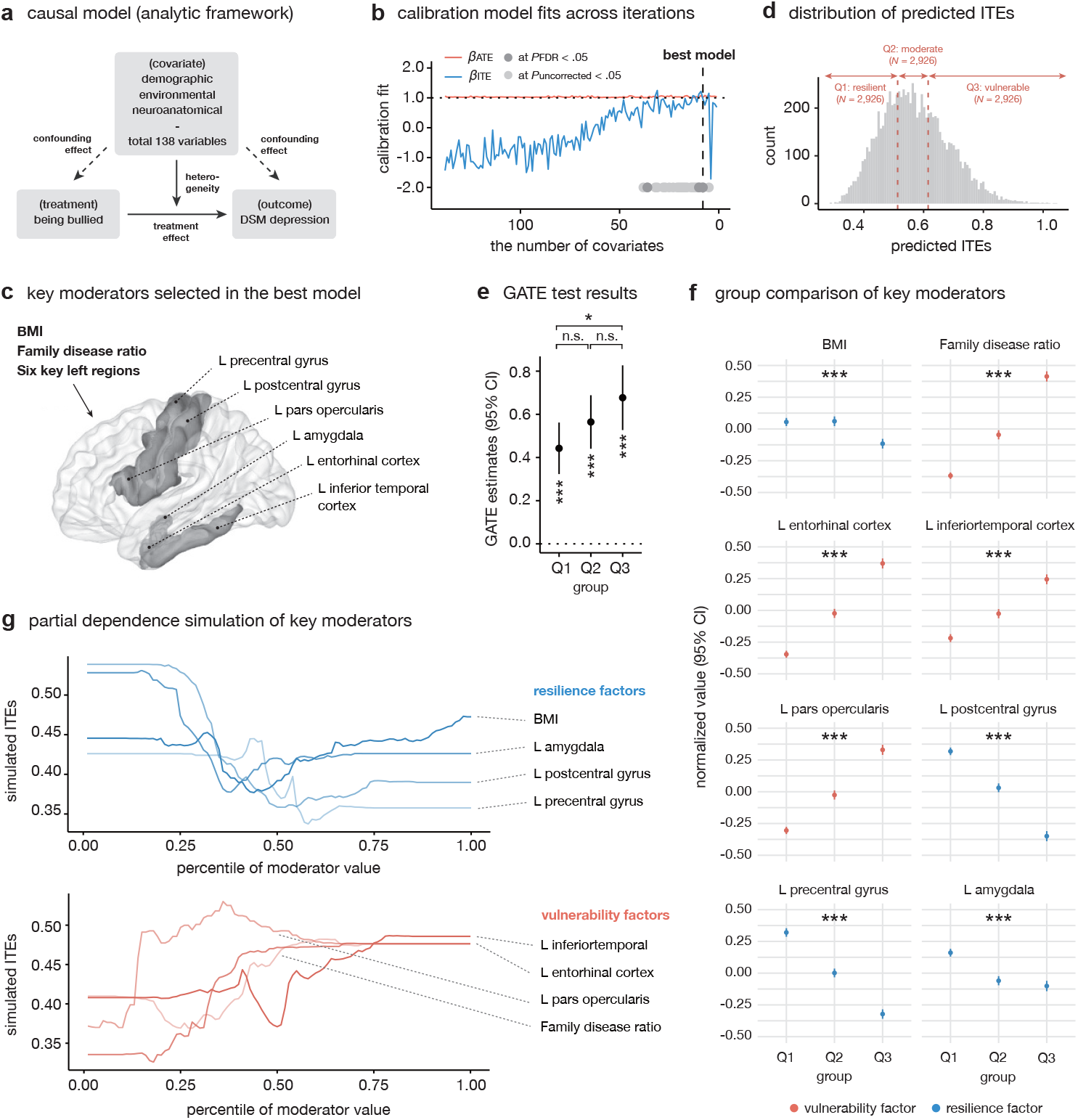
Results of real-world dataset analysis. **a**, Causal model used in real-world analysis. **b**, Calibration model fits across iterations. The vertical dashed line marks the best-calibrated model among 138 models. The horizontal dashed line indicates the calibration test’s gold standard value of 1 (see **Box 2**). **c**, Eight key moderators selected in the best model, visualized through *BrainNetViewer* (Xia et al., 2013). ‘L’ denotes the left hemisphere. **d**, Distribution of predicted ITEs (individualized treatment effects). **e**, GATE (Group-level Average Treatment Effect) results. GATE estimates and pairwise comparisons using two-sided post-hoc *t*-tests. **f**, Group comparisons of eight key moderators. Blue variables indicate resilience factors (significantly lower in higher ITE groups), and red variables indicate vulnerability factors (significantly higher). Statistical differences were examined by one-way ANOVA. **g**, Partial dependence simulations for the eight key moderators. \**P*_FDR_ < .05; \*\**P*_FDR_ < .01; \*\*\**P*_FDR_ < .001.

### Model fitting

We briefly highlight three key considerations when fitting a GRF model in the context of observational behavioral studies: seed ensemble, expected outcome and propensity scores, and clustering variable. See **Appendix Code 1** for model fitting scripts.

#### Seed ensemble

In this tutorial, we build a ‘big model’ consisting of 10,000 trees at each iteration by ensembling five independently trained GRF models, each built using 2,000 trees with a different random seed. Specifically, models from each seed are stored as a list, and once all five are completed, they are merged using the `merge_forests` function. From this ‘big model’, variable importance scores of all covariates are computed using the `variable_importance` function, and the covariate with the lowest importance is excluded from the next iteration. To support model selection, each iteration’s big model is also used to record two key indicators: (1) the minimum and maximum estimated propensity scores (to evaluate the overlap assumption), and (2) the calibration test estimates — β_*ATE*_ and β_*ITE*_.

#### Expected outcomes and propensity scores

When training a GRF model, the `Y.hat` and `W.hat` arguments are used to provide each sample’s expected outcome and propensity score based on covariate profiles — *E*[*Y*_*i*_|*X*_*i*_] and *E*[*W*_*i*_|*X*_*i*_], respectively. These quantities are crucial for predicting ITEs, as they guide how much weight each comparison sample receives based on similarity in covariate profile. In randomized controlled trials (RCTs), where treatment assignment is fully random and independent of covariates, `W.hat` is typically set to 0.5. In contrast, in observational studies that include most psychological and neuroscientific research, both `Y.hat` and `W.hat` should be left as `NULL`. In that case, GRF automatically predicts these quantities internally based on the honesty tree principle (Athey et al., 2019; see **Box 1**).

#### Clustering variable

In the model fitting process, it is important to consider the use of a clustering variable especially in observational studies. GRF assumes that all samples are independently and identically distributed in the covariate space (i.e., *i*.*i*.*d*. assumption). However, this assumption is often violated in psychological and neuroscientific research. For instance, children from the same geographic region may share similar covariate and treatment profiles, leading to correlated outcomes.

To account for such cluster-level confounding, researchers can use clustering variables such as residential site information (e.g., Spechler et al., 2023). Technically, this involves modifying the unit of subsampling: instead of randomly selecting individual samples to build each tree, entire clusters (e.g., sites) are sampled. This prevents data leakage and enforces the honesty property of the forest. That is, the model uses samples from one cluster for training and different clusters for inference, thereby ensuring that training data does not leak into prediction. This design improves the generalizability of model predictions to unseen clusters, such as new data collection sites, not included in training (Athey & Wager, 2019).

Then, how should one decide whether to include a clustering variable, and which one to use? One practical strategy is to run a one-way ANOVA to test whether the outcome variable varies significantly across candidate clustering factors. In our dataset, the outcome variable — CBCL *t*-scores of depressive symptoms significantly differed by data collection site (*F*(21, 8756) = 7.328, *P* < 2×10^-16^). Therefore, we use this site variable as the clustering factor in model fitting throughout the tutorial.

### Model selection

Through the iterative process described above, we fit a total 138 ‘big models’ across iterations. Based on the two criteria (i.e., the overlap assumption and calibration test) and fit index, we identified the model with eight covariates as the ‘best model’ (see **Fig 4b**). The best model chose BMI, family disease ratio, and six left gray matter volume features (see **Fig 4c**). This suggests that individual differences in the impact of bullying victimization on one’s depressive symptoms might be strongly explained by nonlinear combination of familial and neurobiological profiles. In practice, it is possible that no models satisfy both criteria. When this occurs, it may indicate limited heterogeneity and/or a misspecified causal model. **Box 3** outlines the possible reasons for such problems — depending on which assumption is violated — and provides specific recommendations for improving the analytic design. See **Appendix Code 2** for model selection scripts.

### Heterogeneity assessment

Beyond estimating the ATE at the population level, several methods exist to assess whether the predicted ITEs exhibit meaningful heterogeneity. Two representative approaches include the Group Average Treatment Effect (GATE; Chernozhukov et al., 2018; Jacob, 2019) and the Rank-weighted Average Treatment Effect (RATE; Yadlowsky et al., 2025). In this section, we first estimate the ATE and then focus on the conventional GATE framework to assess ITE heterogeneity. See **Appendix Code 3** for heterogeneity assessment processes.

#### Statistical testing for ATE

The GRF package provides the `average_treatment_effect` function to estimate the ATE along with its standard error. In our example, bullying victimization significantly elevated depressive symptoms, indicating a strong treatment effect (ATE = .561, 95% CI = [.462, .661], *P* < 2×10^-16^).

#### Heterogeneity assessment with predicted ITEs

The GATE test proceeds by first generating out-of-bag predictions of ITEs for each individual using the trained GRF model (see **Box 1**). Based on these predicted values, the sample is divided into subgroups, and the group-level ATEs (i.e., GATE) is estimated within each group. In our tutorial, individuals are split into three groups based on the predicted ITEs of bullying on their depressive symptom scores: those with the lowest predicted ITEs (Q1) are labeled the ‘resilient’ group, those in the middle (Q2) as ‘moderate,’ and those with the highest ITEs (Q3) as the ‘vulnerable’ group (see **Fig 4d**). Afterward, group-level comparisons are then conducted using post hoc *t*-tests to determine whether the GATEs differ significantly across groups. If no significant differences are found, this may suggest that treatment effects might be relatively homogeneous across individuals.

In our example (see **Fig 4e**), all three groups exhibited significantly positive GATEs, implying the significant effect of bullying experiences on depressive symptoms across all groups (Q1: GATE = .443, 95% CI = [.323, .562], *P*_FDR_ = 4×10^-13^; Q2: GATE = .565, 95% CI = [.441, .689], *P*_FDR_ < 2×10^-16^; Q3: GATE = .677, 95% CI = [.527, .827], *P*_FDR_ < 2×10^-16^). Among these, Q3 had a significantly higher GATE than Q1, suggesting the meaningful heterogeneity in treatment effects (mean difference = .234, 95% CI = [.043, .426], *P*_FDR_ = .048). However, the differences between Q2 and Q1, as well as Q3 and Q2, did not reach statistical significance (Q2-Q1: mean difference = .122, 95% CI = [-.050, .294], *P*_FDR_ = .242; Q3-Q2: mean difference = .112, 95% CI = [-.082, .307], *P*_FDR_ = .252).

One may question whether differences in GATE across ITE-based subgroups are merely tautological (i.e., whether such group differences are inevitable simply because individuals were grouped based on predicted ITEs). However, this is not the case. To understand why, it is crucial to recognize that the group-level mean of predicted ITEs is not the same as the corresponding GATE estimate. Predicted ITEs are noisy, difficult-to-estimate quantities, generated using an out-of-bag prediction approach that excludes each sample from the trees used to predict its own treatment effect. In contrast, GATE is calculated as a population-level summary and can be more reliably estimated. Because the two quantities are computed in fundamentally different ways, discrepancies in their relative magnitudes across subgroups are common. For instance, it is entirely possible for the middle group (Q2) to have a higher GATE than the high-ITE group (Q3). Such inconsistencies may arise for a variety of reasons, including poor calibration of the GRF model’s ITE predictions, insufficient sample size within each group for reliable GATE estimation. In such cases, researchers should consider examining the model’s calibration test results or reducing the number of groups to increase statistical power.

### Moderator Identification

Through the analyses above, we observed that the impact of childhood bullying on children’s depressive symptoms shows substantial individual variability, and that this heterogeneity is closely linked to familial and neurobiological profiles. But which specific variables are associated with increases or decreases in ITEs? In this section, we introduce three complementary approaches to explore this question. See **Appendix Code 4** for moderator identification processes.

#### Group comparison

The simplest approach to investigate the relationship between covariates and ITEs is to test whether the groups identified through the GATE test differ significantly on each covariate. We performed one-way ANOVAs to test whether the volume of each region differed across three subgroups. As a result, all eight key moderators showed significant group differences at *P*_FDR_ < .05 (see **Appendix C** for detailed statistics). Among these, four variables showed increasing trends from Q1 to Q3, suggesting their role as ‘vulnerability’ factors, while the other four features showed decreasing ones, suggesting potential ‘resilience’ factors (see **Fig 4f**).

#### Best linear projection

While the group comparison method investigates bivariate associations between each covariate and ITEs, the best linear projection approach takes a multivariate perspective. Using the `best_linear_projection` function in the grf package, we fit a multiple linear regression model in which all eight key moderators were included as regressors to explain variance in predicted ITEs. This method estimates the beta coefficients of each moderator. In our analysis, only family disease ratio showed a statistically significant positive association with ITEs at *P* < .05. The remaining seven moderators did not exhibit significant predictive effects (see **Appendix C** for details).

#### Partial dependence simulation

Although the eight key moderators were selected by the best model as the strongest predictors of ITE heterogeneity, the limited explanatory power observed in the best linear projection suggests a need to account for nonlinear relationship between moderators and ITEs while controlling interactions among them. To address this, we used partial dependence simulation, which offers a model-agnostic framework for examining the marginal effects of individual moderators after controlling for all others.

To probe the role of individual moderators, we simulated ‘hypothetical participants’ in which one target variable was systematically varied while all other variables were held constant. In practice, we fixed the remaining seven covariates at their median values and divided the observed range of the target moderator into 100 percentiles. For each percentile, we generated a simulated sample and entered it into the trained best model to predict ITEs. Repeating this procedure for each of the eight moderators produced curves showing how predicted effects changed as the moderator increased or decreased, while interactions with other covariates were controlled.

These simulations revealed clear nonlinear patterns for most moderators (see **Fig 4g**). For example, BMI initially appeared to act as a resilience factor in the group-comparison analysis, but the simulation uncovered a U-shaped relationship: both low and high BMI predicted stronger treatment effects. This pattern is consistent with prior evidence linking BMI and stress resilience (He et al., 2022; Kim et al., 2015; Viner et al., 2006).

### Summary

In this section, we demonstrate how GRF can be applied to real-world data analysis. Using a backward-elimination model selection procedure, we fit a GRF model that effectively explained the relationship between observed bullying exposure and depressive symptoms. The model not only revealed a strong ATE of bullying exposure but also uncovered substantial individual differences in the ITEs. Among a range of demographic, environmental, and neuroanatomical covariates, familial psychopathologic history, BMI, and gray matter volumes of six regions in the left hemisphere emerged as key moderators most strongly associated with the variance in ITEs. These regions spanned the amygdala, precentral gyrus, entorhinal, inferior temporal and inferior frontal cortices. Importantly, partial dependence simulation revealed that the relationship between these moderators and predicted ITEs was not merely linear, but instead reflected complex, nonlinear associations. This finding opens avenues for generating and testing and replicating new hypotheses about the role of relatively underreported brain regions (e.g., precentral gyrus) in modulating children’s stress responses to early-life adversity. Furthermore, it holds potential for early identification of at-risk individuals and the development of targeted preventive interventions based on neurobiological markers.

## Discussion

In this paper, we introduced a computational framework based on GRF to systematically dissect individual differences in treatment/exposure effects. Our work makes two methodological contributions that significantly enhance the reliability and scope of heterogeneity analysis in psychology and the behavioral sciences: the *seed ensemble* and a *backward elimination* approach for model selection.

The first contribution, the seed-ensemble method, addresses a key vulnerability in causal machine learning: sensitivity of results to algorithmic stochasticity. Models trained with different random seeds can produce conflicting conclusions about the existence of heterogeneity, undermining reproducibility. In psychiatry and related fields, where such analyses can inform diagnosis and treatment, this instability is unacceptable. By aggregating models trained under multiple seeds, our ensemble approach yields more stable ITE predictions. Simulations further showed that this method improves reliability more effectively than simply increasing the number of trees within a single-seed model, improving the inferential integrity of GRF analyses.

The second contribution provides a principled solution to model selection in high-dimensional spaces. Many neurocognitive functions and psychopathologies arise from complex, distributed mechanisms, which require comprehensive, data-driven frameworks such as whole-brain modeling (Borsboom et al., 2018; Krakauer et al., 2017; Menon et al., 2023; Olthof et al., 2023). Our backward elimination framework meets this need by systematically pruning covariates while enforcing calibration between estimated latent treatment effects and observed dataset. Rather than producing a final definitive model, our framework functions as a powerful engine for hypothesis generation, isolating a parsimonious set of moderators that best explain heterogeneity. In our simulation analysis, this approach demonstrated superior identification performance across various scenarios compared to the conventional heuristic, which interprets variables with the top *n*% of variable importance from a fully fitted model using all covariates as key moderators. Considering that our approach does not require the arbitrary specification of the importance threshold *n*, it exhibits high accuracy and adaptability in most research settings where the true number of key moderators is unknown. Furthermore, our approach maintained solid performance not only on the train set used for model selection but also on the test set, suggesting that it provides generalizable interpretations rather than being overfitted to a given dataset.

Particularly, our application of the framework to the ABCD dataset demonstrates how GRF provides knowledge-aligned and novel findings regarding key factors of individual differences. Among the six primary gray matter volume variables selected through a backward-elimination process, the left amygdala, postcentral gyrus, precentral gyrus, and inferior temporal gyrus exhibited a U-shaped relationship with simulated ITEs. This finding aligns with nonlinear models describing the relationship between the structure of specific regions and their functions. That is, gray matter volumes that are either too small or too large may imply insufficient or excessive synaptic pruning and myelination during child development (Glasser & Van Essen, 2011; Gogtay et al., 2004; Kassem et al., 2013; Whitaker et al., 2016). It has been consistently reported that when core emotional processing regions like the amygdala show such structural variations, the adverse mental health effects of stress are amplified (Jones et al., 2019; Liu et al., 2024; Weissman et al., 2020). The same U-shaped pattern observed between BMI and ITEs can also be explained by the interplay between chronic inflammation from excess body fat and the nutritional imbalances and depression vulnerability associated with insufficient body fat (Byrne et al., 2015; de Wit et al., 2009; Kim et al., 2015; McLachlan et al., 2023; Moazzami et al., 2019). This demonstrates that our GRF framework can accurately capture the nonlinear relationships between specific treatment effects and their key moderators.

However, some of our findings are relatively novel, highlighting GRF’s potential for generating new, data-driven hypotheses. For instance, while some studies have suggested that the postcentral and precentral gyri encode affective afferents such as pain and emotional expressions (Cao et al., 2018; Vogt, 2005), their specific role in modulating social stress responses in conjunction with other regions has not been clearly established. One hypothesis is that their engagement in interoceptive attention, performed in concert with subcortical regions like the amygdala, may be crucial in moderating the affective outcomes of stressful events. Indeed, the primary motor and somatosensory cortices are closely involved in perceiving bodily responses to threatening stimuli (Critchley et al., 2004; Khalsa et al., 2009; Pollatos et al., 2007; Terasawa et al., 2013), and such efficient interoceptive awareness can serve as a foundation for effective emotion regulation (Fermin et al., 2024; Haase et al., 2016). To test this hypothesis, further investigation is needed to determine if these regions are identified as key moderators in new datasets, and more controlled experiments are required to verify their causal relationships.

We note that our approach reduces the need for domain knowledge, not eliminating them. The validity of our causal claims rests on the untestable assumption of unconfoundedness that all common confounders have been included in the covariate set. Indiscriminately including all available variables in a model risks introducing collider bias, creating spurious associations (Akimova et al., 2021; Elwert & Winship, 2014). Thus, thoughtful, theory-informed covariate selection, perhaps guided by causal graph theory, should precede data-driven analyses. Also, it should be noted that our implementation focuses on the canonical Causal Forest algorithm with a binary treatment. The GRF ecosystem offers additional tools, including Instrumental Forests to address unobserved confounding, Quantile Forests for distributional effects, and methods for continuous treatments. These extensions represent promising directions for future work. The principles of our framework, however, are broadly applicable and can be extended to various data modalities (e.g., functional connectivity, genetics) and research questions across the behavioral and brain sciences.

### Limitations

Despite its strengths, our framework has several limitations that suggest avenues for future research. First, the variable importance metrics that guide backward elimination-based model selection approach can be biased when covariates are highly correlated each other, potentially overstating the importance of certain variables while underrepresenting others (Strobl et al., 2008; Tolosi & Lengauer, 2011). Incorporating conditional importance metrics (Strobl et al., 2008) or permutation-based alternatives (Gong et al., 2021) may help mitigate these biases. Second, the variable importance calculation in random forest systematically underestimates the importance of categorical variables (Hines et al., 2022; Strobl et al., 2007). Bias-corrected split criteria (Hothorn et al., 2006) and importance metrics (Hines et al., 2022) could offer more balanced variable importance estimates. Last, the iterative nature of the backward elimination process is computationally intensive, particularly for datasets with thousands of features. Future work could focus on developing more computationally efficient feature selection algorithms.

## Conclusion

This work demonstrates how GRF can be adapted to move beyond population averages toward a person-centered science of behavior by predicting individualized treatment effects and identifying key moderators of these personalized estimates. By devising and validating analytic approaches that enable GRF to perform these tasks reliably even with high-dimensional datasets, we demonstrated that the GRF framework can generate more comprehensive hypotheses about individual difference factors and thereby facilitate subsequent research. Furthermore, by data-driven identification of responsive subgroups to specific treatments, we highlighted its potential to inform the development of more efficient and personalized clinical and policy interventions. In doing so, it contributes to a broader shift in psychology and behavioral sciences: treating heterogeneity not as unexplained noise, but as a central feature of human functioning and opportunity for evidence-based personalization.

## Acknowledgement

This work was supported by the National Research Foundation of Korea (NRF) grant funded by the Korea government (MSIT)(No. 2021R1C1C1006503, RS-2023-00266787, RS-2023-00265406, RS-2024-00421268, RS-2024-00342301, RS-2024-00435727, NRF-2021M3E5D2A01022515), by Creative-Pioneering Researchers Program through Seoul National University (No. 200-20240057, 200-20240135), by Semi-Supervised Learning Research Grant by SAMSUNG (No.A0342-20220009), by Identify the network of brain preparation steps for concentration Research Grant by LooxidLabs (No.339-20230001), by Institute of Information & communications Technology Planning & Evaluation (IITP) grant funded by the Korea government (MSIT) [NO.RS-2021-II211343, 2021-0-01343, Artificial Intelligence Graduate School Program (Seoul National University)] by the MSIT (Ministry of Science, ICT), Korea, under the Global Research Support Program in the Digital Field program (RS-2024-00421268) supervised by the IITP (Institute for Information & Communications Technology Planning & Evaluation), by the National Supercomputing Center with supercomputing resources including technical support (KSC-2023-CRE-0568), by the Ministry of Education of the Republic of Korea and the National Research Foundation of Korea (NRF-2021S1A3A2A02090597), by the Korea Health Industry Development Institute (KHIDI), and by the Ministry of Health and Welfare, Republic of Korea (HR22C1605), by Artificial intelligence industrial convergence cluster development project funded by the Ministry of Science and ICT (MSIT, Korea) & Gwangju Metropolitan City and by KBRI basic research program through Korea Brain Research Institute funded by Ministry of Science and ICT (25-BR-05-01).

## Author Contributions

**Jinwoo Lee:** conceptualization, analysis, visualization, writing, review, editing, code arrangement.

**Junghoon Justin Park:** conceptualization, analysis, writing, review.

**Maria Pak:** conceptualization, analysis, visualization, writing, review.

**Seung Yun Choi:** conceptualization, analysis, writing, review.

**Jiook Cha:** conceptualization, writing, review, funding acquisition.

## Box 1 Honesty rule of GRF to prevent data leakage without explicit train/test split

GRF uses an ‘honest’ subsampling design to ensure a rigorous inference of average and individual treatment effects: each tree’s subsample is split into two disjoint parts — one half is used to determine the tree’s splitting structure, and the other half is used to estimate leaf-level parameters (e.g., leaf-level average treatment effects). This separation between structure-learning and parameter-estimation eliminates the bias introduced by adaptively fitting splits.

In practice, the honesty mechanism functions much like having separate training, validation, and test sets within each tree: one part defines the tree structure, another estimates the parameters, and out-of-bag (OOB) samples — those unused in both steps — serve as implicit test data. For example, when predicting sample *i*’s ITE, τ_*i*_, GRF uses only those trees where sample *i* was not involved in either structure learning or parameter estimation (i.e., truly “honest” trees). This yields OOB predictions without the need for explicit cross-validation or train/test splits, preserving estimation integrity and enabling rigorous inference for all samples.

The figure below shows how GRF applies the honesty rule when building trees and how out-of-bag (OOB) predictions for ITEs are obtained. For illustration, consider a toy example with eight samples and four trees. In each tree, three samples are randomly chosen for splitting the tree structure (group A, shown in blue), and another three distinct samples are chosen for estimating treatment effects within the leaves (group B, shown in red). The remaining two samples are unused in that tree (shown in gray).

To predict the ITE for sample ①, GRF only uses trees in which sample ① was not involved in either splitting or estimation. In this example, Tree 3 meets that condition. Within Tree 3, sample ① is dropped down the tree, and any group B samples (e.g., ④, ⑦, or ③) that land in the same terminal node are considered similar and assigned weights. By aggregating these weighted neighbors across unrelated trees, GRF obtains an out-of-bag estimate of the ITE for each sample. This mechanism allows GRF to make predictions for all units without requiring an explicit train/test split.

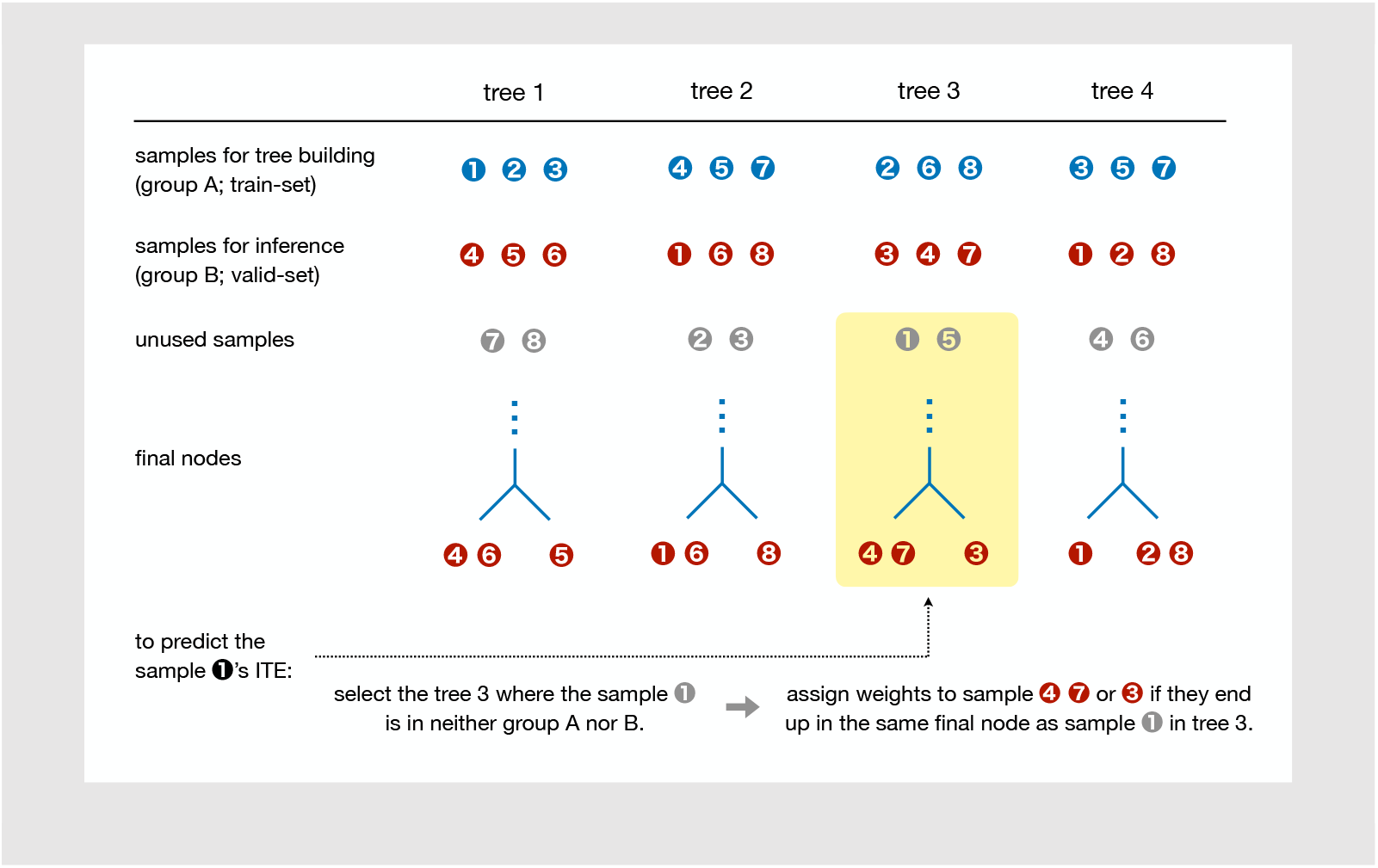

## Box 2 Calibration test

The calibration test is a type of omnibus test used to evaluate the fit of a GRF model. It assesses whether the predicted ATE and ITEs accurately capture the relationship among covariates, treatment, and outcomes in the observed data. The degree of calibration for the ATE and ITE is evaluated using the following equation:

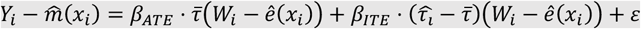

In this equation, *Y*_*i*_ is the observed outcome of the sample *i*,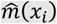 is the expected outcome given covariates 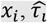 is the predicted ITE of the sample *i*, and 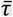 is the average of predicted ITEs across all samples (i.e., predicted ATE). *W*_*i*_ is the observed treatment, and *ê*(*x*_*i*_) is the predicted propensity score.

Conceptually, this regression evaluates whether the residual variation in outcomes after adjusting for predicted outcomes can be explained by the model’s prediction of ATE and ITE. Specifically, the calibration coefficients *β*_*ATE*_ and *β*_*ITE*_ quantify how well the predicted ATE and individual-level variation in ITE align with the observed data. When both coefficients are statistically close to 1, this indicates that the model’s treatment effect predictions are well-calibrated to the structure of the observed outcome and treatment data.

It is important to interpret these metrics with caution. Unlike typical predictive performance metrics in conventional machine learning, the calibration coefficients do not directly indicate whether the ATE or ITE “accurately” predicted, since the true values of these causal quantities are unobservable (i.e., the fundamental problem of causal inference). Rather, the calibration test assesses whether the predicted ATE and ITE explain the observable data in a statistically consistent way. It evaluates internal coherence, not ground-truth validity.

## Box 3 The absence of the ‘best’ model – why and how?

### Case 1: When the overlap assumption failed

A violation of the overlap assumption typically arises when there is severe imbalance between treated and untreated samples in the dataset. The ITE for a given individual is calculated based on comparisons with individuals sharing similar covariate profiles who received the treatment and those who did not. However, when the data are heavily skewed (e.g., if untreated individuals vastly outnumber treated ones), it becomes hard to find treated samples with comparable covariate profiles. In such cases, researchers should first check for sample imbalance and consider corrective approaches such as up-or down-sampling to restore comparability across groups.

### Case 2: When the calibration test failed

On the other hand, it is possible that none of the models trained through the backward elimination process pass the calibration test. Specifically, this occurs when the estimates of β_*ATE*_ and/or β_*ITE*_ are not statistically significant at *P*_FDR_ < .05 in any of the models. Since *β*_*ATE*_ and *β*_*ITE*_ reflect how well the predicted ATE and ITE capture the latent structure of the observed treatment and outcome data, a failure to meet this threshold suggests that the model’s predictions are not well aligned with the actual data (see **Box 2**).

In many cases, predicting individual-level ITEs is substantially more challenging than calculating ATE, so it is common for *β*_*ITE*_ to be non-significant. If *β*_*ATE*_ is significant while *β*_*ITE*_ is not, this may indicate that the GRF model successfully captured the population-level ATE but failed to accurately predict sample-level ITEs. This could occur if the treatment variable chosen by the researcher has little or no causal impact on the outcome, or if key moderators that shape the distribution of treatment effects are missing from the covariate set.

To address this issue, one possible solution is to consider alternative treatment variables that may be more causally relevant to the outcome. Additionally, expanding the covariate set guided by domain expertise to include theoretically meaningful moderators may improve the model’s calibration. It is also important to note that these problems may simply reflect insufficient sample size to support stable estimation of both ATE and ITE.

### Appendix Methods 1. Seed Ensemble Simulation

#### A. Data Simulation

To validate our proposed solutions, we generated partially simulated datasets, combining real-world covariates with simulated treatment and outcome variables. This approach is advantageous because it preserves the complex and realistic data structures (e.g., multicollinearity) of empirical data while still allowing for a known ground truth treatment effect. This method is an alternative to using purely artificial data, which can often lack realism (Jacob, 2021). A similar strategy has been effectively used in healthcare GRF research, where real patient covariates were used to simulate outcomes (Wendling et al., 2018). Adopting this framework, we used real covariates from a large-scale neuroimaging study and simulated the treatment assignment and outcome variables.

##### A-(1). Covariates

Covariates were sourced from the Adolescent Brain Cognitive Development (ABCD) Study Data Release 5.1. We selected demographic covariates, including the child’s age, sex, race, family income, parental educational level, parental marital status, and study site. Neuroimaging covariates consisted of structural MRI (sMRI) brain volume metrics from cortical and subcortical segmentations (Desikan et al., 2006; Fischl et al., 2002). Following the ABCD study’s recommendations, we excluded 1,174 subjects who did not meet the sMRI inclusion criteria (Casey et al., 2018). For the remaining 9,623 subjects, we preprocessed the brain variables by normalizing them by total intracranial volume and removing any near-zero-variance features. To isolate the contributions of specific regional brain structures, we also excluded global measures (e.g., total hemispheric volumes) and ventricular volumes. From this processed dataset, we randomly selected a subsample of 1,000 subjects to ensure computational feasibility. Nominal covariates were converted into dummy variables, and all continuous variables were z-scaled. The final set of covariates contained 1,000 subjects and 149 features with no missing values.

##### A-(2). ITE simulation

Adapting the simulation algorithm from Jacob (2021), we generated four simulation datasets based on a 2×2 design, varying the linearity/nonlinearity of the ITE generating function and the strength (strong/weak) of the heterogeneity. To enhance the ecological validity of our simulation and ensure its relevance to common research scenarios in neuroscience, we deliberately selected heterogeneity-driving variables based on prior empirical findings rather than using arbitrary ones. Specifically, drawing upon the findings of Dick et al. (2021), who demonstrated that pre-existing neural features in specific threat-processing regions moderate stress-related outcomes, we selected the volumes of the right amygdala, right anterior cingulate cortex, and left parahippocampal gyrus as the key variables (*x*_1_, *x*_2_, *x*_3_) expected to drive treatment effect heterogeneity. This approach allows us to test our framework’s performance in a scenario that mimics plausible, empirically-supported neurobiological moderation effects.

To simulate an observational study containing confounding between treatment and outcome, we generated the binary treatment variable *W* via a nonrandom assignment mechanism. A key challenge in such studies is that the factors influencing treatment assignment are often also related to the outcome. To operationalize this principle in our simulation, we modeled the propensity score, *e*(*x*), as a logistic function of the first two heterogeneity variables. The treatment *W* was then drawn from a Bernoulli distribution based on this propensity score:

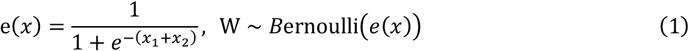

The individual treatment effect (ITE), τ, was then simulated in two forms: a linear and a nonlinear version, both dependent on the three key moderators (*x*_1_, *x*_2_, *x*_3_). The equations are as follows:

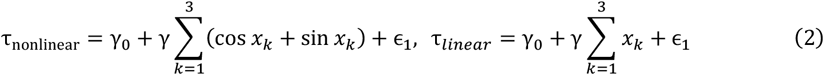

Here, γ_0_ and γ are tunable parameters used to control the magnitude of the treatment effect, and ϵ_1_ is a random Gaussian noise term. Notably, in the linear condition, this formulation means that each of the three heterogeneity variables (*x*_1_, *x*_2_, *x*_3_) was set to contribute equally to the overall treatment effect, as they share a common coefficient, γ. Finally, the outcome *Y* was simulated using a partially linear model incorporating the main effects of the heterogeneity variables, the treatment status *W*and the treatment effect τ:

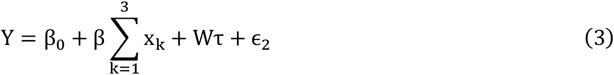

In this model, β_0_ and β are tunable parameters, and ϵ_2_ is a random Gaussian noise term. Our goal was to create two distinct conditions (“weak” and “strong” heterogeneity) for both the linear and nonlinear scenarios. The magnitude of this ground-truth heterogeneity is primarily controlled by the γ parameter in the ITE equation.

However, rather than defining our conditions by the value of γ itself, we defined them based on the statistical power of a GRF model to detect the resulting heterogeneity. To achieve this, we employed an iterative tuning process: we systematically adjusted the tunable parameters (primarily γ_0_ and γ) and, for each parameter set, generated a full dataset and trained a corresponding GRF model. We then used the `test_calibration` function to obtain the *P*-value for β_*ITE*_. This process was repeated until we identified parameter sets that consistently produced *P*-values within our target ranges. This resulted in the following operational definitions for our two conditions:

- **Weak heterogeneity:** Datasets generated using parameters that yielded a *P*-value for β_*ITE*_ in the range 0.05 < *P* ≤ 10^™5.^
- **Strong heterogeneity:** Datasets generated using parameters that yielded a *P*-value for β_*ITE*_ of *P* ≤ 10^™5^.

#### B. Analysis

##### B-(1). Comparison of calibration metrics between single-seed and seed-ensembled models

We first qualitatively compared the reliability of calibration metrics (i.e., β_ATE_ and *β*_ITE_) between single-seed models and seed-ensembled models, while controlling for the total number of trees (see **Fig 2a** and **2b**). This analysis used the dataset generated under conditions of weak heterogeneity and a nonlinear ITE function (see *‘A. Data Simulation’* above). We calculated the calibration metrics for four single-seed models (1,000 trees each) and four seed-ensembled models (200 trees fitted on 5 different seeds). This design ensured that four models were fitted per condition, each with a total of 1,000 trees. A total of 20 random seeds were sampled without replacement and organized into four groups of five seeds. The ensembled models were fitted using all five seeds within their respective group, whereas the single-seed models were fitted using only the first seed from each group. Finally, we qualitatively assessed the variance of *β*_ATE_ and *β*_ITE_ based on their point estimates and 95% confidence intervals among the four models within each condition.

##### B-(2). Extended grid analysis

Next, to quantitatively evaluate these differences across a wider range of datasets and scenarios, we conducted a grid analysis (see **Fig 2c** and **Appendix A**). This analysis utilized all four simulated datasets, extending beyond the weak heterogeneity and nonlinear ITE condition. For each dataset, we fitted 50 models per condition using different the number of tree and seed combinations, spanning a grid of 16 conditions (seed counts: 1, 2, 5, 10; tree counts: 1,000, 2,000, 5,000, 10,000). For each of the 50 models within a given condition, we computed the negative log-transformed *P*-values for *β*_ATE_ and *β*_ITE_. We then calculated the coefficient of variance of these values across the 50 models to quantify the metric’s variance for that specific condition.

Finally, we assessed not only the calibration metrics but also the validity and reliability of the GRF model’s ITE predictions against the true simulated ITE, as a function of seed and tree combinations (see **Fig 2d**). For a fair comparison, this analysis focused on four conditions, all maintaining a total of 10,000 trees: (n_seeds, n_trees) pairs of (1, 10,000), (2, 5,000), (5, 2,000), and (10, 1,000). For each of the 50 models fitted per condition, we calculated Pearson’s *r* between the predicted ITE and the true ITE as the prediction performance metric. A one-way ANOVA was conducted to test for significant differences in the mean prediction performance across the four conditions. Furthermore, Levene’s test was performed to assess significant differences in the variance of the prediction performance. All *P*-values were FDR-corrected for each statistical testing.

### Appendix Methods 2. Backward Elimination Simulation

#### A. Data Simulation

##### A-(1). Train set for model selection

We utilized the weak heterogeneity datasets under both linear and nonlinear conditions to conduct backward-elimination model selection and evaluate whether this approach can accurately identify key moderators that modulate the simulated ITE. For details on the simulation process for these two datasets, please refer to Section *‘A. Data Simulation’* in **Appendix Methods 1**.

##### A-(2). Test set for generalizability testing

Furthermore, to assess the generalizability of ‘the best model’ selected via the backward elimination process, we evaluated its ability to accurately predict ITE not only on the train set also on an independent set. To this end, we generated new, independent test sets for both the linear and nonlinear conditions, respectively. In these independent sets, we generated the noise terms ϵ_1_ and ϵ_2_ in the equation (2) and (3) using different random seeds, compared to the train sets mentioned above. The noise magnitude (standard deviation) and the rest of the algorithm stayed the same as in the train sets.

#### B. Analysis

To empirically validate our proposed backward elimination-based model selection framework, we tested its ability to identify true key moderators that drive heterogeneity of ITEs. We compared the performance of our method against a common heuristic approach using the two simulated datasets: one with linear and another with nonlinear treatment effects, both designed to have weak heterogeneity. The strong heterogeneity conditions were intentionally excluded from this evaluation; the effect-modifying variables in those datasets were so prominent that nearly any selection method could identify them, thus providing limited discriminative power for comparing framework performance.

##### B-(1). Backward elimination-based model selection

For both the linear and nonlinear datasets, we first implemented our backward elimination procedure. The initial step involved training a full GRF model that included all 149 candidate covariates. To ensure a robust and stable initial model, we used a seed ensemble approach, constructing the model by merging five individual forests of 2,000 trees each, with each forest trained using a different random seed.

Following the training of this full ensemble model, we initiated the iterative backward elimination process. In each iteration, the variable with the lowest importance score, as determined by the `variable_importance` function on the current model, was removed. This process was repeated until only one covariate remained, generating a sequence of 149 candidate models of decreasing complexity. This entire sequence of models was then subjected to our two-stage selection criteria. First, we retained only those models that passed the calibration tests for both β_*ATE*_ and β_*ITE*_ at a FDR–corrected significance level of *P* < .05 and that also satisfied the overlap assumption (i.e., all predicted propensity scores *ê* falling within the [0.05, 0.95] interval). From this filtered set of valid models, we selected the single best model as the one that minimized our specified fit index:

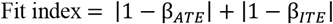

##### B-(2). Baseline – “top 10%” heuristic

For comparison, we evaluated a common heuristic method for variable selection. This involved training a single, full GRF seed ensemble model on all 149 covariates, using the identical configuration as in our method (i.e., an ensemble of 5 forests of 2,000 trees each). From this model, we extracted the variable importance scores and selected the top 10% of covariates, resulting in a fixed set of 15 variables.

##### B-(3). Comparison of identification and prediction performance

First, the moderator identification performance of both our backward elimination framework and the heuristic method was quantified by their F1 scores. Considering the large class imbalance in this task (i.e., choosing three variables from 149 ones), we chose the F1 score as a performance metric that assesses precision and recall simultaneously (see **Fig 3a**). Second, we evaluated the ITE prediction performance of ‘the best model’ selected via backward-elimination model-selection process on both the train and test sets, under linear and nonlinear conditions, respectively. The train set was directly used for model selection. For the test set, ITEs for the test samples were directly inferred without any additional training, using the `‘newdata’` argument of the `‘predict’` function. The predictive performance metric was calculated as the Pearson’s *r* between the true ITE and the predicted ITE (see **Fig 3b**).

### Appendix Methods 3. Real-world Dataset: ABCD study

The real-world data tutorial was conducted using baseline data from the Adolescent Brain Cognitive Development (ABCD) Study, Release 5.1. The analysis aimed to identify heterogeneity in the effect of bullying experience (treatment) on the Child Behavior Checklist (CBCL)(Achenbach, 2011) DSM depressive symptoms *t*-score (outcome). The initial dataset included a total of 138 covariates, encompassing demographic factors (e.g., age, biological sex, race/ethnicity), environmental variables (e.g., caregiver’s income, marital status, and education level, the rate of psychiatric disorders among family members), and neurobiological features comprising body mass index and gray matter volume metrics across whole-brain cortical and subcortical regions. See **Appendix B** for participants’ demographic and environmental profiles.

This raw data underwent a thorough preprocessing pipeline for analysis. First, samples with any missing values were excluded from the analyses. Second, based on inspection of variable distributions, two samples with undefined factors in the biological sex column and 433 outliers in body mass index (defined as values exceeding 1.5 times the interquartile range) were removed. Third, all gray matter volume metrics were normalized by dividing them by whole-brain volume. Fourth, we confirmed that none of the variables exhibited near zero-variance. Finally, continuous variables were z-transformed, and nominal variables were one-hot encoded, including the first factor level. This pipeline yielded a final, clean dataset prepared for the GRF analysis framework.

## Appendix A. Reliability gain of seed ensemble in simulation analyses

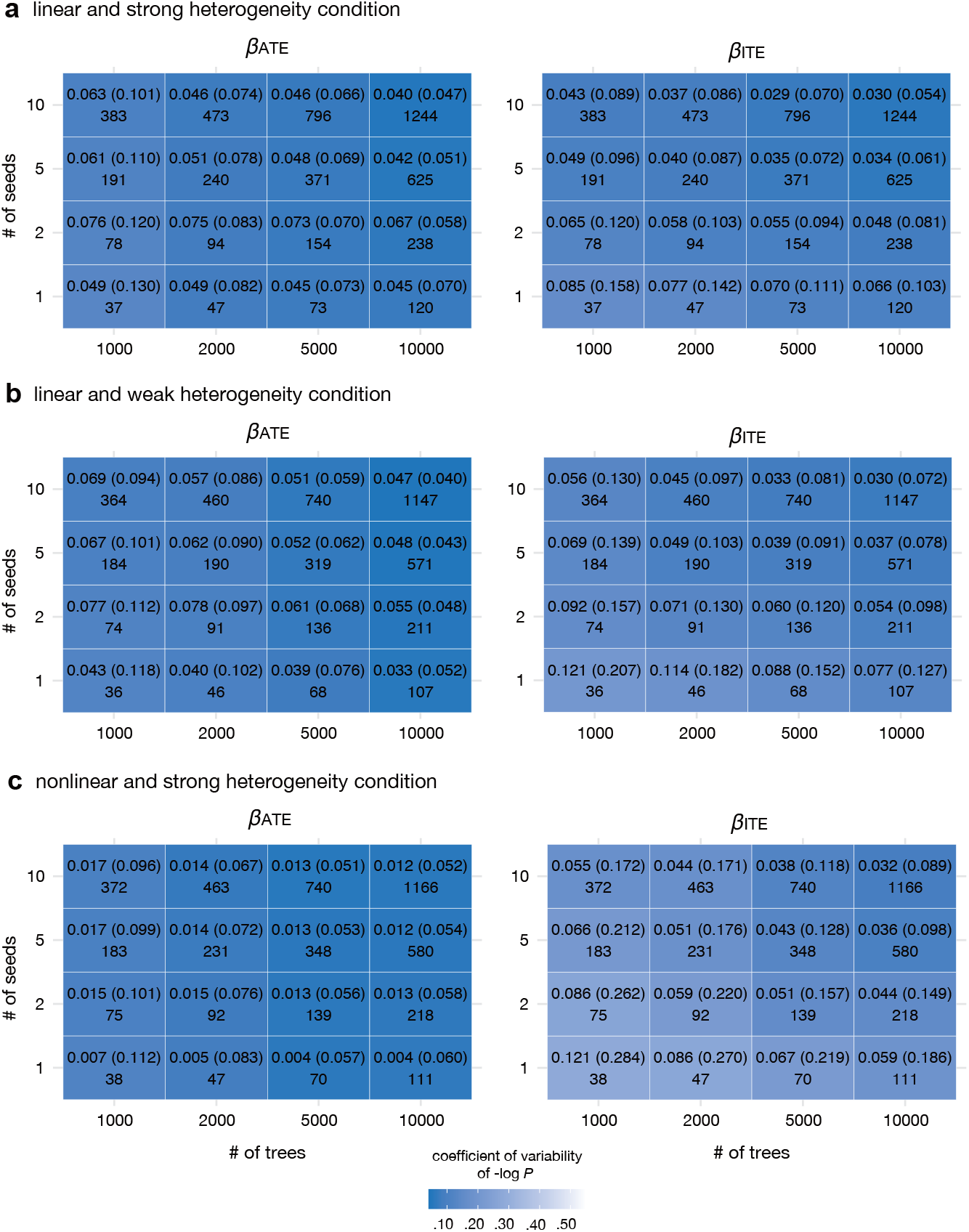

We examined the reliability gain of calibration metrics from seed ensemble approach across four simulation conditions defined by two factors: 1) whether each sample’s ITE follows a linear or nonlinear function of treatment and key moderators, and 2) whether the degree of heterogeneity is strong or weak (see **Appendix Methods 1**). Each cell in the resulting grid corresponds to a (tree, seed) configuration, reporting the coefficient of variation of the estimated beta coefficients and their –log *P*-values across 50 trials, as well as the total computation time in seconds. Results for the nonlinear and weak heterogeneity condition, most representative of real-world datasets, are presented in **Fig 2c** of the main text.

Under linear heterogeneity, both β_*ATE*_ and β_*ITE*_ showed low variability even with a single seed and a small number of trees (see **panels a** and **b**). In contrast, under nonlinear conditions, regardless of heterogeneity strength, using a seed ensemble was more effective than merely increasing the number of trees in reducing the variability of *β*_*ATE*_ and *β*_*ITE*_ estimates and their *P*-values, which supports the utility of seed ensemble to achieve more seed-consistent model fittings in a real-world dataset analysis (see **panel c** and **Fig 2c** in the main text).

## Appendix B. Demographics and environmental profiles in real-world tutorial

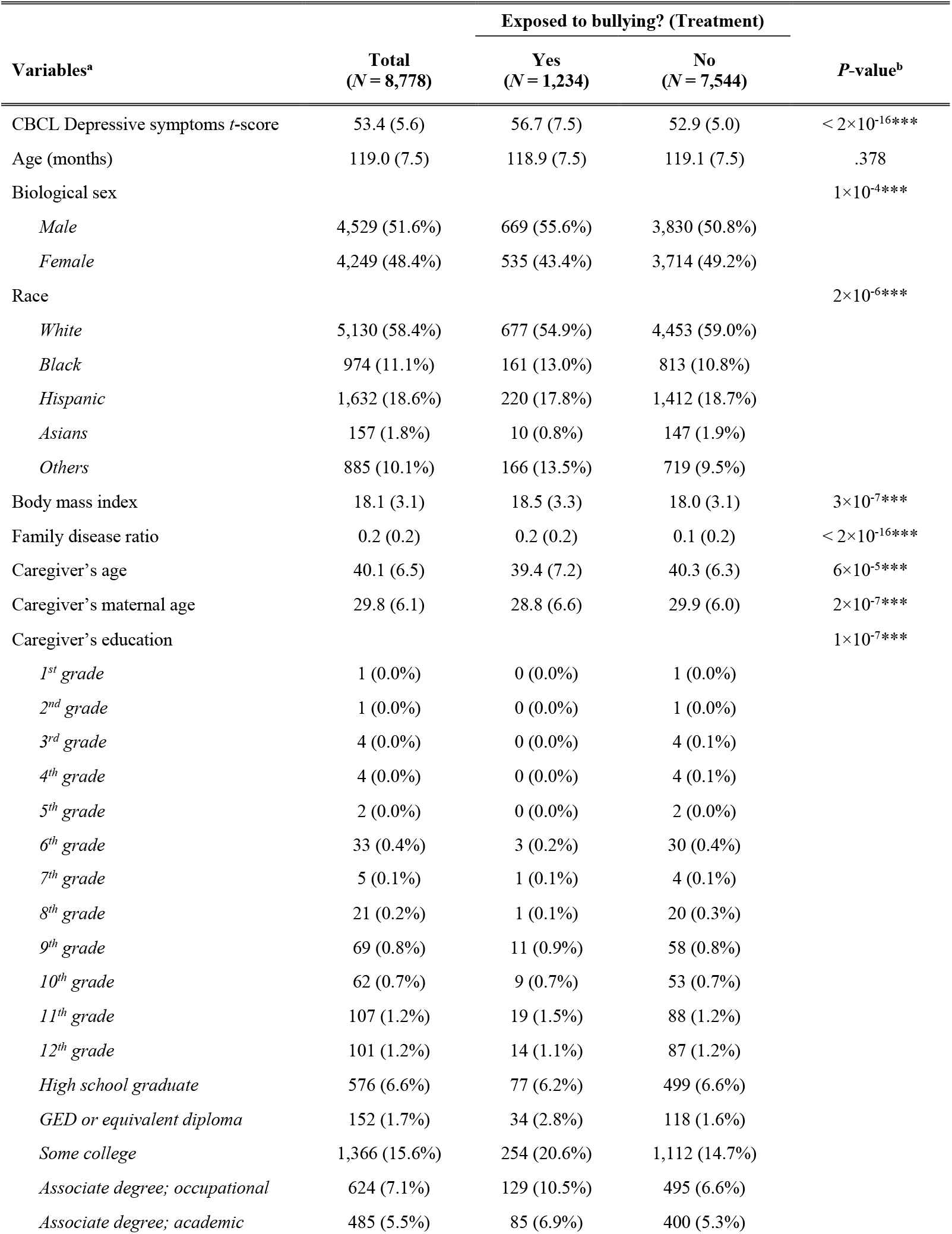
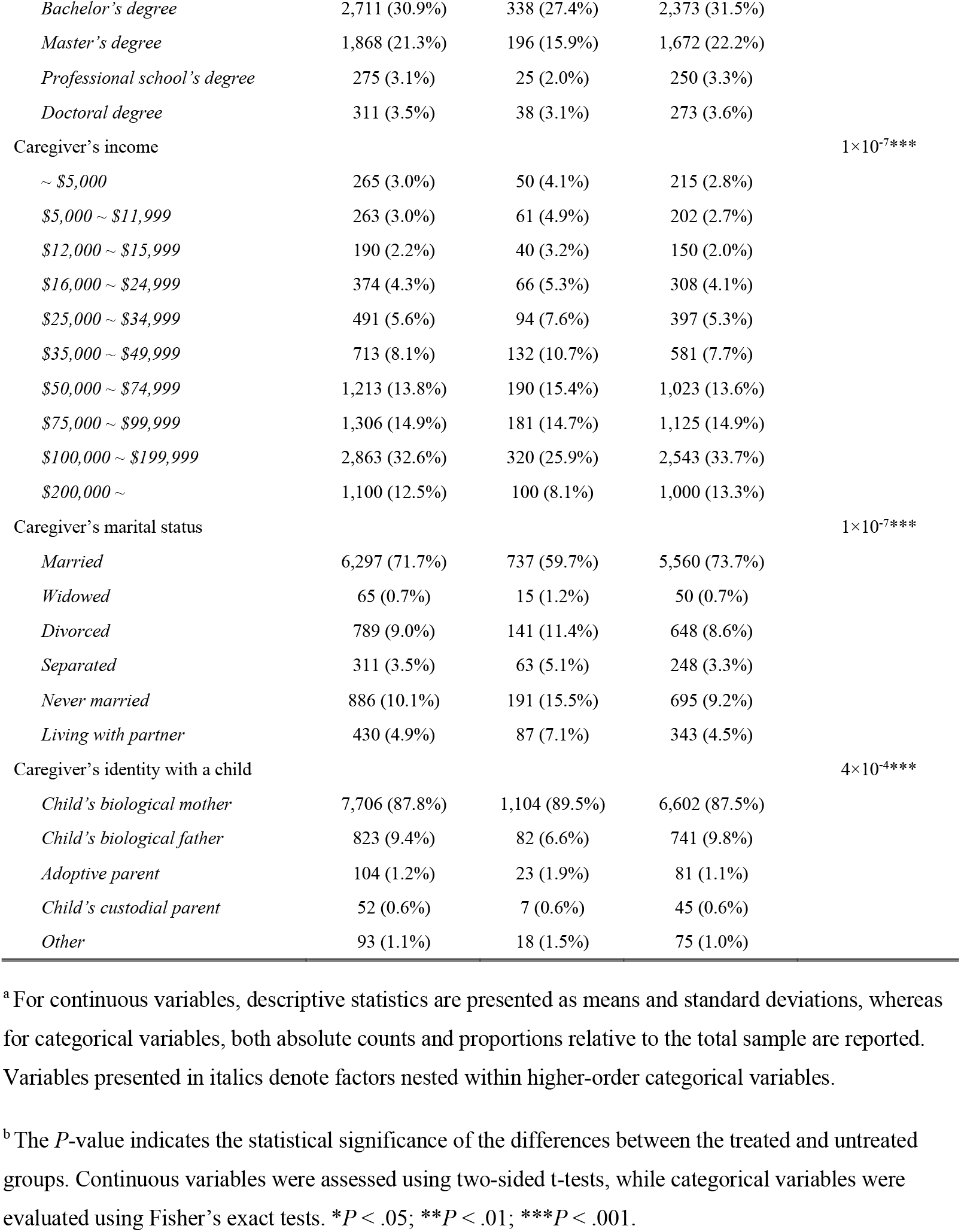

## Appendix C. Statistics of selected key eight covariates in moderator identification analyses

| Selected moderators | Group comparison |  |  |  |  | Best linear projection |  |  |
| --- | --- | --- | --- | --- | --- | --- | --- | --- |
| | Q1<br>*M(SD)* | Q2<br>*M(SD)* | Q3<br>*M(SD)* | *F*(2, 8775) | *P*<sub>FDR</sub> | $\beta$ | 95% CI | *P* |
| BMI | .055<br>(.881) | .061<br>(1.050) | -.116<br>(1.050) | 29.624 | $2 \times 10^{-13}***$ | -.021 | [-.116, .075] | .671 |
| Family disease ratio | -.368<br>(.697) | -.046<br>(.991) | .414<br>(1.107) | 502.974 | $< 2 \times 10^{-16}***$ | .091 | [.025, .157] | .007** |
| L entorhinal cortex | -.346<br>(.771) | -.024<br>(.981) | .369<br>(1.090) | 410.412 | $< 2 \times 10^{-16}***$ | .078 | [-.026, .181] | .141 |
| L inferior temporal cortex | -.219<br>(.871) | -.026<br>(.995) | .245<br>(1.069) | 164.415 | $< 2 \times 10^{-16}***$ | .056 | [-.025, .137] | .174 |
| L pars opercularis | -.305<br>(.785) | -.025<br>(.996) | .330<br>(1.091) | 317.594 | $< 2 \times 10^{-16}***$ | .072 | [-.018, .161] | .117 |
| L postcentral gyrus | .318<br>(.822) | .031<br>(.952) | -.350<br>(1.092) | 354.941 | $< 2 \times 10^{-16}***$ | -.060 | [-.135, .015] | .119 |
| L precentral gyrus | .320<br>(.927) | .002<br>(.948) | -.322<br>(1.018) | 324.182 | $< 2 \times 10^{-16}***$ | -.030 | [-.148, .087] | .615 |
| L amygdala | .161<br>(.847) | -.060<br>(.989) | -.101<br>(1.124) | 58.843 | $< 2 \times 10^{-16}***$ | -.029 | [-.116, .057] | .507 |

In the group comparison analysis, group-wise differences of each covariate were examined using one-way ANOVA, and all *P*-values were FDR-corrected. Best linear projection analysis was conducted using the `best_linear_projection` function in the GRF package. Since a single model was fitted, *P*-values for the beta coefficients of each variable were not multiple-corrected. In the ‘covariate’ column, “L” and denotes the left hemisphere. All variables were z-transformed. \**P* < .05; \*\**P* < .01; \*\*\**P* < .001.

### Appendix Codes

These code snippets are pseudo-code implementations of the analytical framework presented in the “Tutorial with Real-world Dataset” section of the main draft, using the `grf` package in R Studio. They have been edited to help readers understand the structure and logic of the code. For the actual implementation, please refer to the official GitHub repository. This section consists of four connected code blocks corresponding to model fitting, model selection, heterogeneity assessment, and moderator identification. Functions mentioned in the main draft (e.g., `merge_forests`) are highlighted in bold.

#### Appendix Code 1. Model fitting

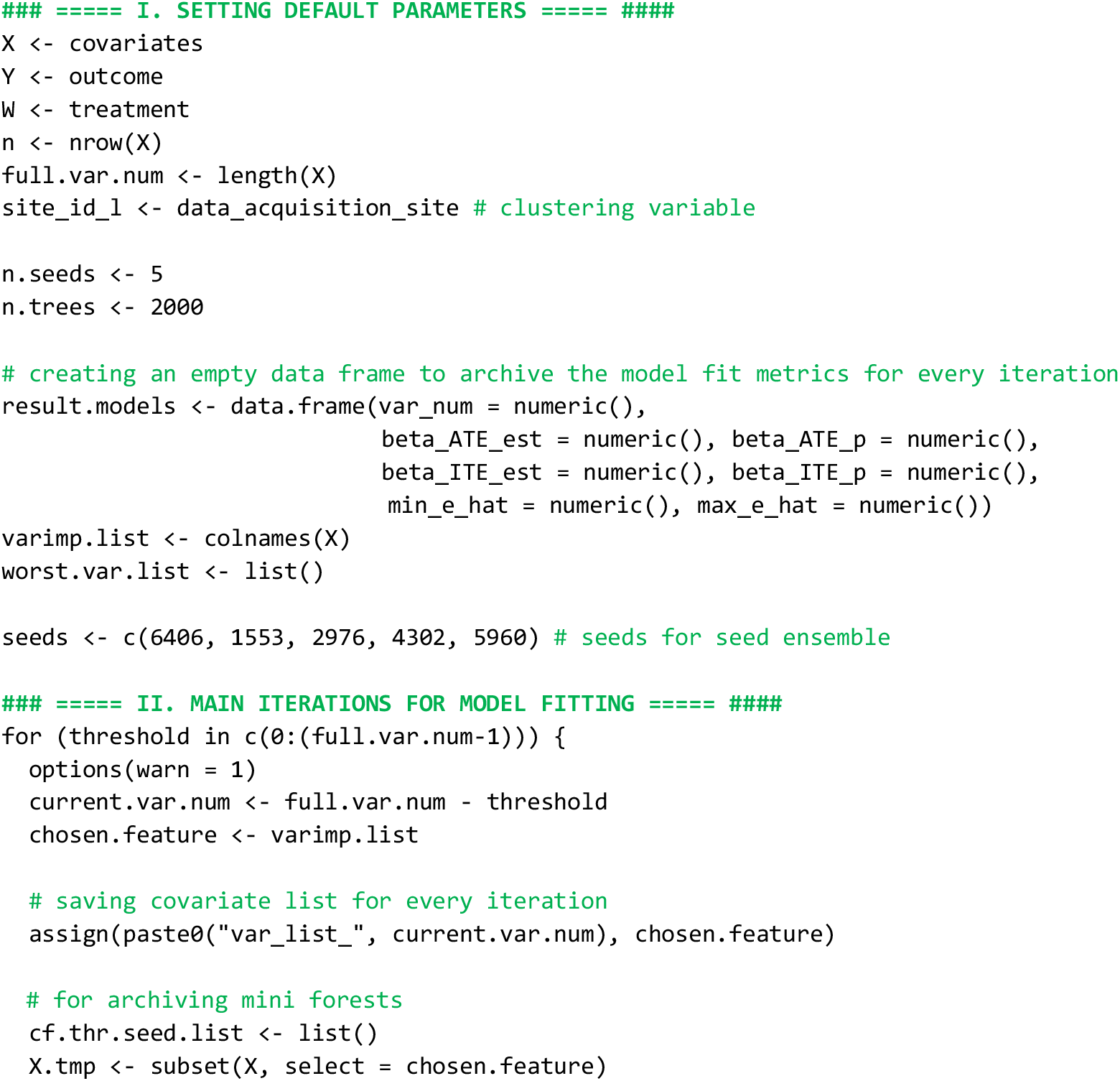
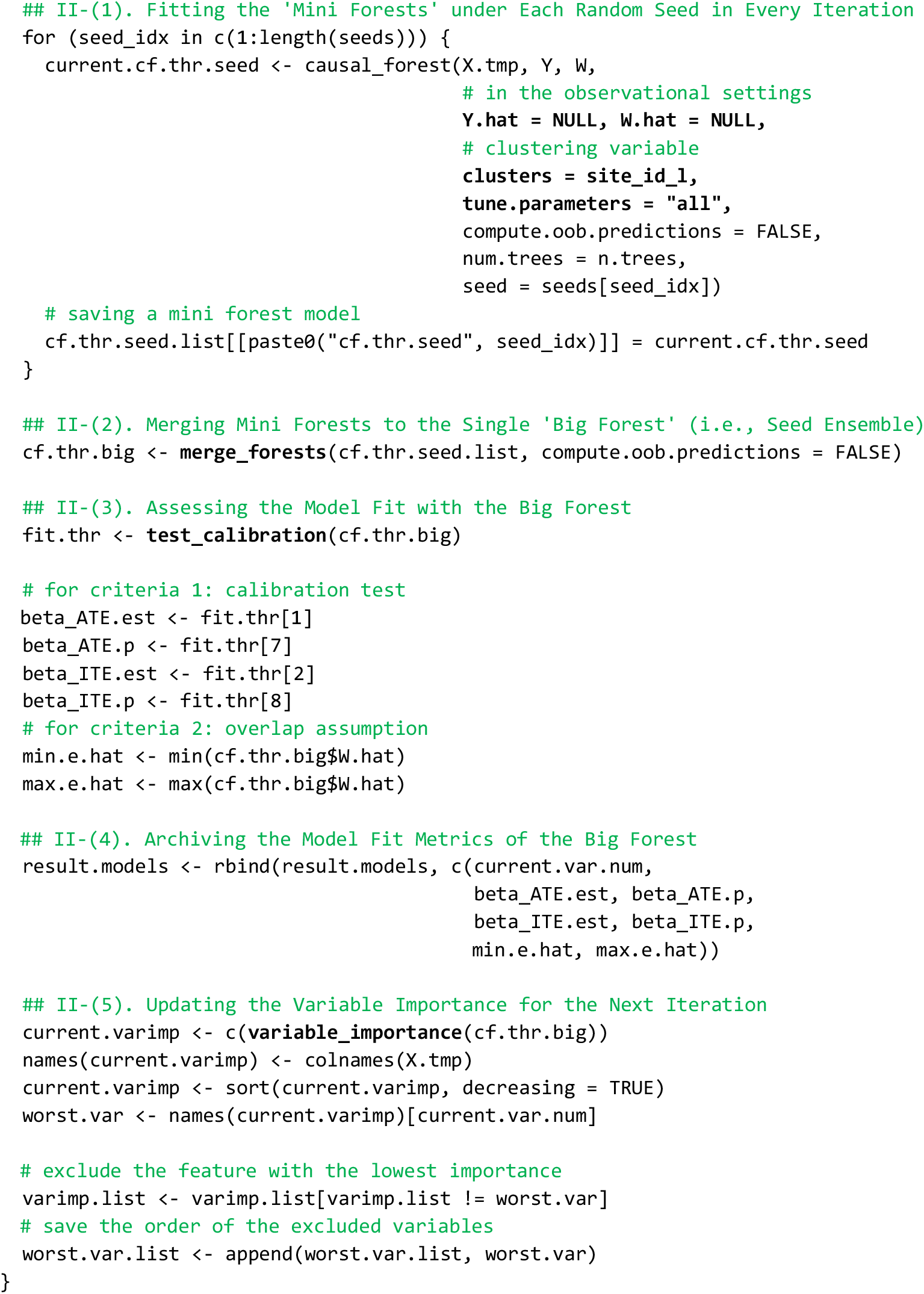

#### Appendix Code 2. Model selection

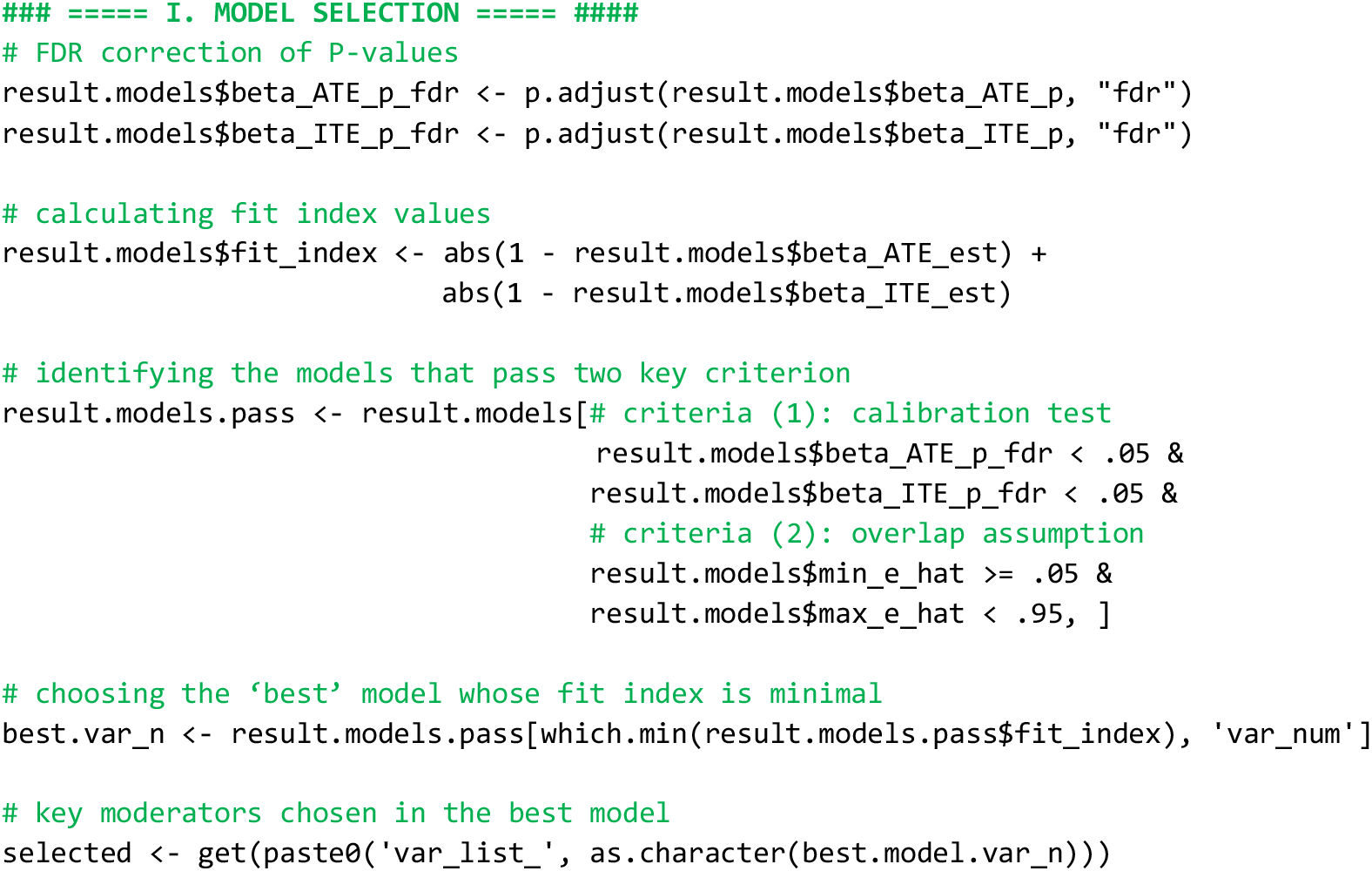

#### Appendix Code 3. Heterogeneity assessment

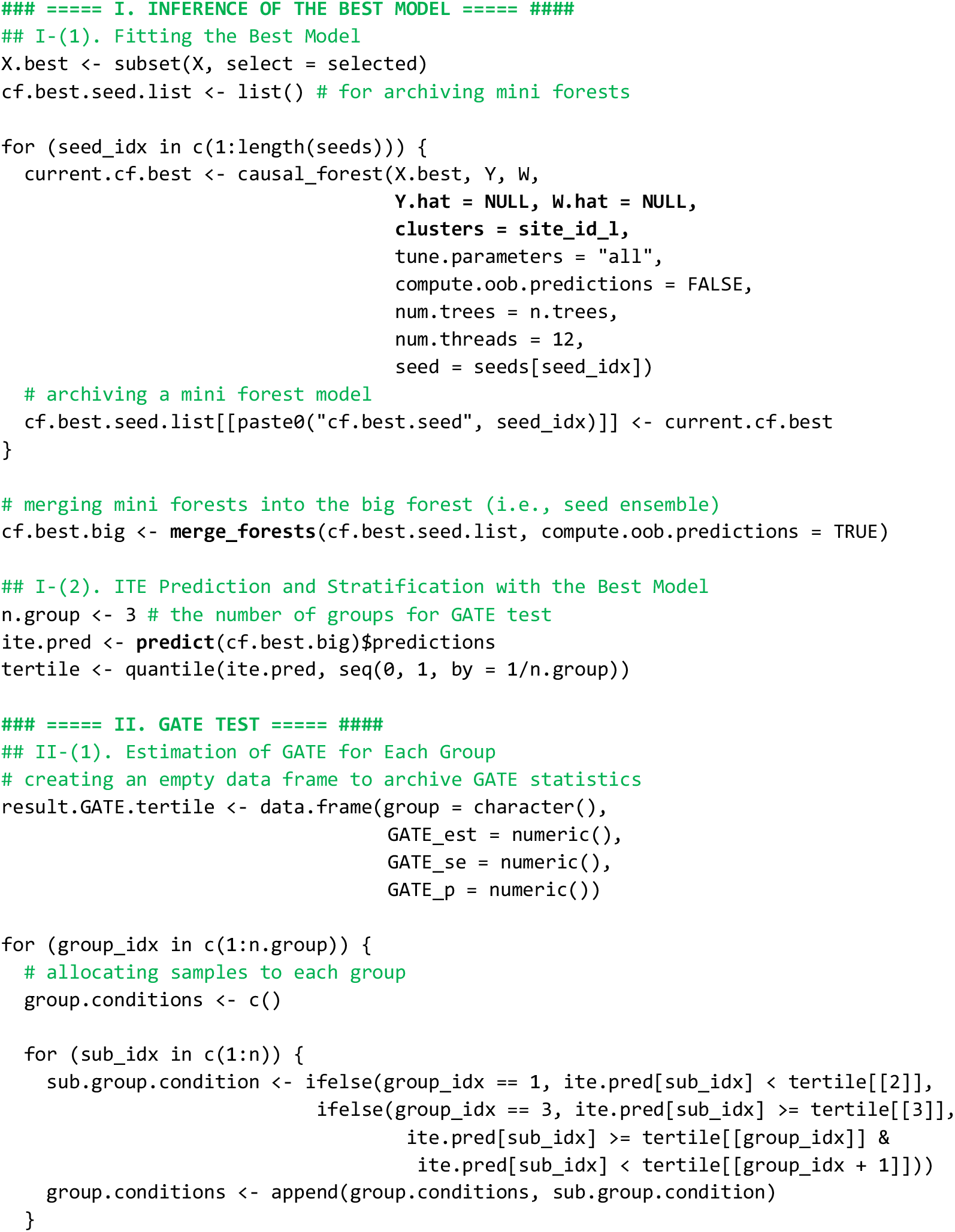
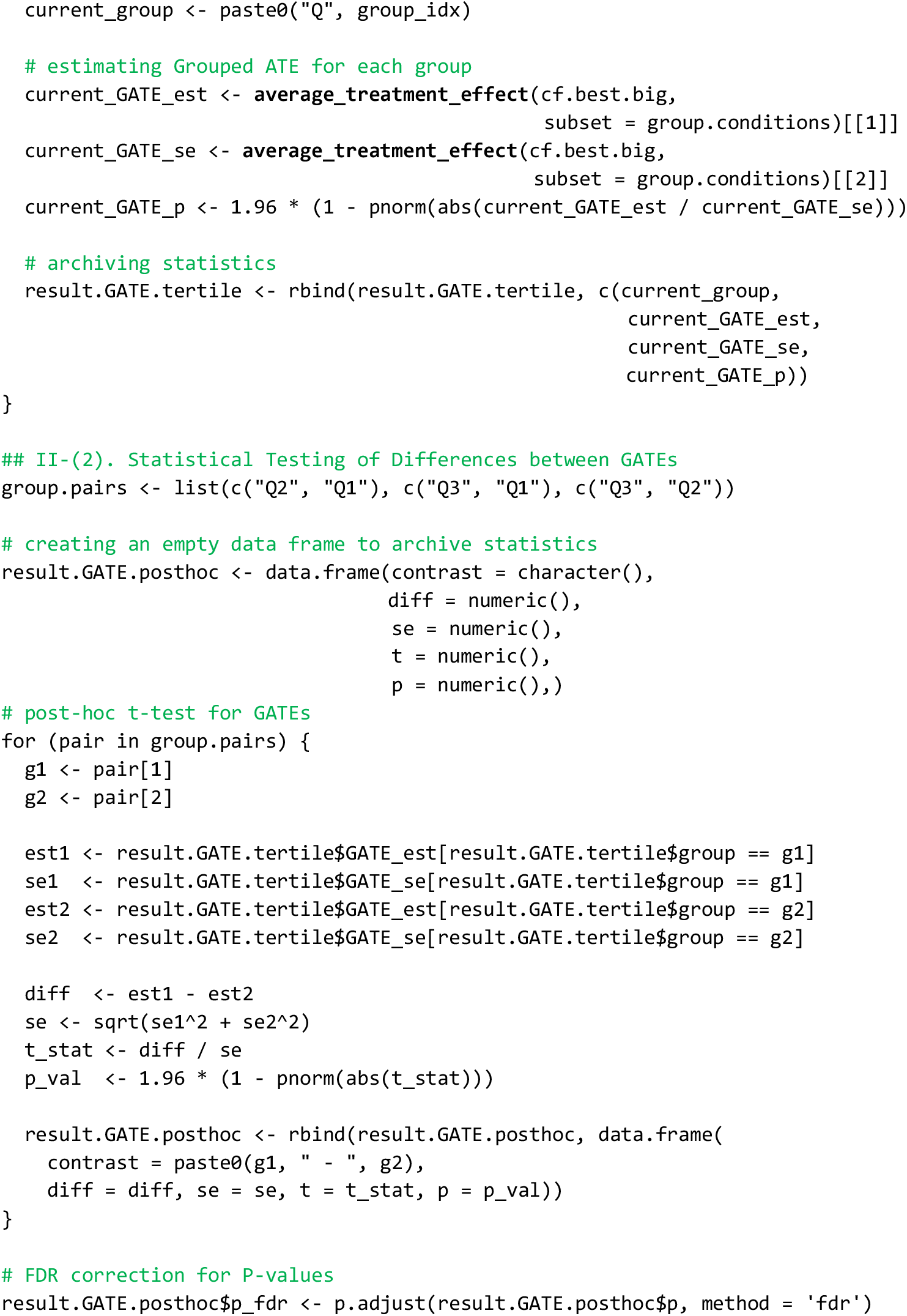

#### Appendix Code 4. Moderator identification

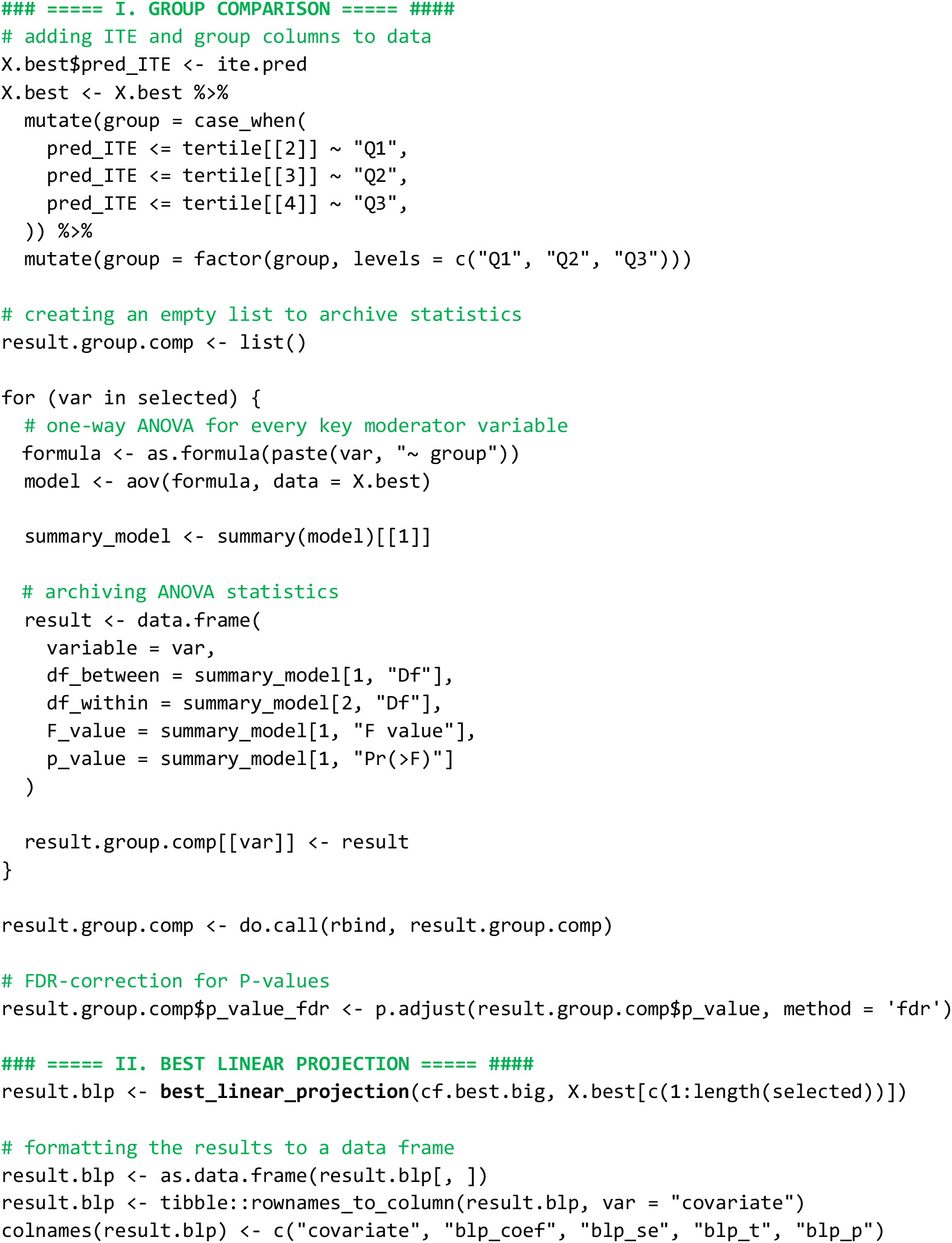
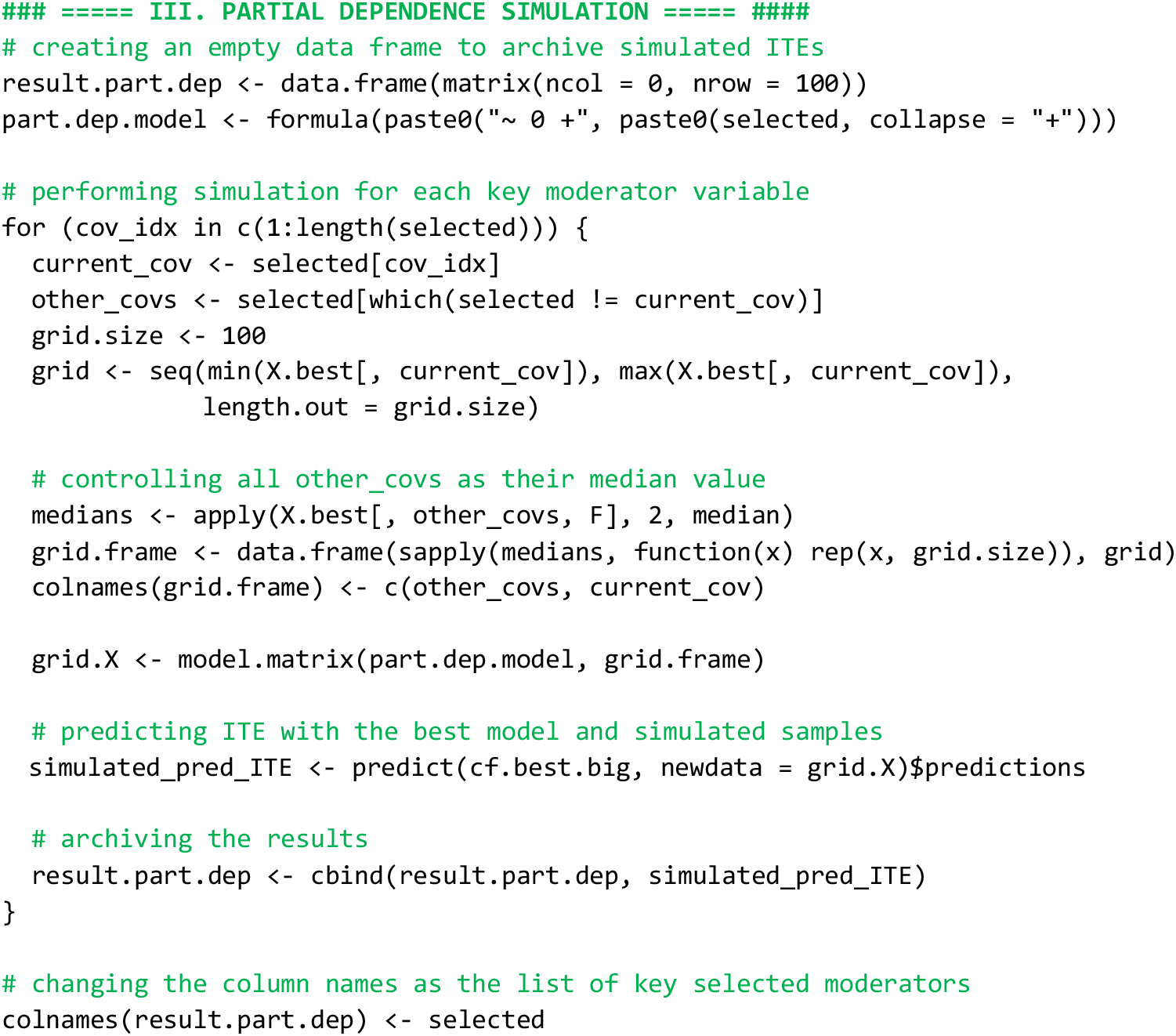

## Footnotes

1 In GRF, the covariate set consists of variables that act as confounders affecting both the cause and the outcome, while also serving as potential candidates for moderating the treatment effect. GRF performs leaf splits using covariates that maximize differences in treatment effects across leaves, thereby learning to both control for these variables and identify key moderators. In this paper, all such variables will be collectively referred to as “covariates” during the model fitting process. Subsequently, in the moderator identification stage based on variable importance, only the key variables will be post hoc labeled as “moderators.”

